# Distinct features of ribonucleotides within genomic DNA in Aicardi-Goutières syndrome (AGS)-ortholog mutants of *Saccharomyces cerevisiae*

**DOI:** 10.1101/2023.10.02.560505

**Authors:** Deepali L. Kundnani, Taehwan Yang, Alli L. Gombolay, Kuntal Mukherjee, Gary Newnam, Chance Meers, Zeel H. Mehta, Celine Mouawad, Francesca Storici

## Abstract

Ribonucleoside monophosphates (rNMPs) are abundantly found within genomic DNA of cells. The embedded rNMPs alter DNA properties and impact genome stability. Mutations in ribonuclease (RNase) H2, a key enzyme for rNMP removal, are associated with the Aicardi-Goutières syndrome (AGS), a severe neurological disorder. Here, we engineered two AGS-ortholog mutations in *Saccharomyces cerevisiae*: *rnh201*-G42S and *rnh203*-K46W. Using the ribose-seq technique and the Ribose-Map bioinformatics toolkit, we unveiled rNMP abundance, composition, hotspots, and sequence context in these yeast AGS-ortholog mutants. We found higher rNMP incorporation in the nuclear genome of *rnh201*-G42S than in wild-type and *rnh203-*K46W-mutant cells, and an elevated rCMP content in both mutants. Moreover, we uncovered unique rNMP patterns in each mutant, highlighting a differential activity of the AGS mutants towards rNMPs embedded on the leading or on the lagging strand of DNA replication. This study guides future research on rNMP characteristics in human genomic samples carrying AGS mutations.

## INTRODUCTION

Aicardi-Goutières syndrome (AGS) is a rare genetic disorder that predominantly affects the brain, skin, and immune system^1^. AGS is typically diagnosed in infancy or early childhood and is characterized by a range of neurological symptoms including developmental delays, intellectual disabilities, seizures, and movement abnormalities^2^. AGS patients present with mutations in nucleic-acid metabolism regulators such as TREX1 (Three Prime Repair Exonuclease 1), SAMHD1 (SAM And HD Domain Containing Deoxynucleoside Triphosphate Triphosphohydrolase 1), RNASEH2A/B/C (Ribonuclease H2 Complex), ADAR (Adenosine Deaminase RNA Specific), and IFIH1 (Interferon Induced with Helicase C Domain 1)^3^. More than 50% of AGS patients exhibit mutations in the RNase H2-complex^4^.

RNase H2 is a critical enzyme complex involved in ribonucleotide excision repair (RER) during DNA replication and repair. The incorporation of ribonucleotides in DNA happens in the form of ribonucleoside monophosphates (rNMPs) and can occur due to various factors, including incorporation by DNA polymerases or embedment of RNA primers during DNA synthesis^3,4^. Failure to efficiently remove ribonucleotides can contribute to disease, compromising the integrity of the genome^5,6^, and rNMPs can hinder the interaction of proteins with DNA during DNA replication and transcription, potentially disrupting normal cellular functions^7^. Therefore, the efficient removal of rNMPs by RNase H2 is vital for maintaining genomic stability and preventing the accumulation of DNA damage^8^.

Human RNase H2 consists of 3 subunits (A, B and C) with RNase H2A as its catalytic subunit. In yeast, the orthologous genes to human RNase H2 Subunit A, B and C are *RNH201*, *RNH202*, and *RNH203*, respectively ^9–11^. Studies in yeast have revealed that mutations in these genes lead to genomic instability, DNA damage, and replication defects^12–14^ . Defective RNase H2 function in yeast increases embedded rNMP presence, leading to replication stress, DNA breaks, and activation of damage response pathways, revealing the detrimental effects of rNMP incorporation and genome instability^12,15^. Previous studies reporting rNMPs in *Saccharomyces cerevisiae* with *rnh201*△ genotype show preference for deoxyadenosine immediately upstream from the most abundant rCMPs and rGMPs in the nuclear genome^16^. Frequencies and patterns of rNMP incorporation around early firing autonomously replicating sequences (ARSs) in *rnh201*△ yeast strains also show distinct upstream nucleotide preferences for deoxyadenosine in the leading strand, and deoxycytidine in the lagging strand^17^. Moreover, several reports have shown that RNase H2 is not active in yeast and human mitochondrial DNA^18,19^ and imbalances in the total cellular dNTP pool led to alteration in the frequency of rNMPs incorporated into mtDNA in yeast^16,18^.

Out of the mutations reported in RNase H2 enzyme complex, the G37S mutation occurring close to the catalytic site residues of the RNase H2 subunit A protein (highlight with green circles in Figure 1), has been extensively studied^10^ . G37S mutation in RNase H2A leads to reduced catalytic activity of RNase H2, invokes cGAS–STING innate immune-sensing pathway, and is perinatally lethal in mice^20^. RNase H2A subunit is also seen to be well conserved among eukaryotic species in contrast to RNaseH2B and RNase H2C (Figure 1). In the enzyme structural complex, RNaseH2C subunit is closer to the catalytic site of subunit A in the ribonuclease complex^10^, and among AGS mutations in RNase H2B and C, the R69W mutation in RNaseH2C has been observed to significantly affect the rNMP-cleavage specificity, highlighting its role in enzymatic function^21^. Moreover, the human RNASEH2A-G37S and RNASEH2C- R69W mutations of RNase H2 are present in regions that are conserved in the yeast *Saccharomyces cerevisiae* ortholog proteins Rnh201 and Rnh203, respectively (Figure 1). Here, we engineered yeast strains with the human RNASEH2A-G37S and RNASEH2C-R69W orthologs mutations in the *RNH201* and *RNH203* genes, respectively, of the yeast BY4742 background (*MAT*α *his3*Δ1 *leu2*Δ0 *lys2*Δ0 *ura3*Δ0) to study rNMP incorporation patterns in these RNase H2 mutants.

**Figure 1.**
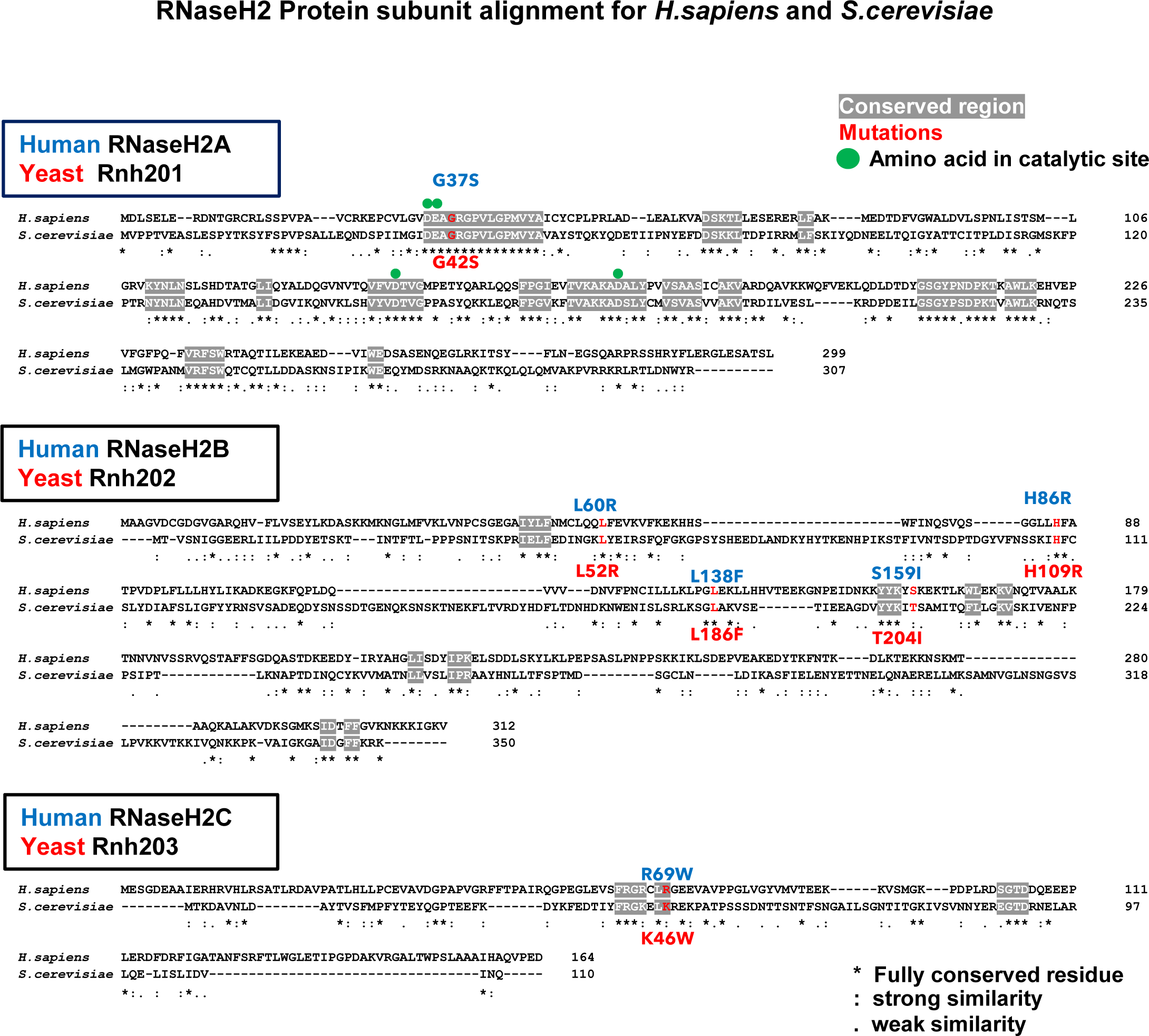
Human and Yeast RNase H2 subunit amino-acid sequence alignment and AGS mutations commonly studied. Alignment of amino acid sequences of RNase H2 catalytic subunit H2A, and accessory subunits H2B and H2C from *H. sapiens* and *S. cerevisiae*^32^. The grey highlights show the most conserved parts of the protein complex^32^. Amino acids highlighted in red show AGS mutations in human RNase H2 subunits and the corresponding mutations in the *S. cerevisiae* orthologs enzyme subunits within fully conserved or similar amino acids^47^. The amino acids with green circle on top represent those present in the catalytic site of the enzyme subunit A^9^. The mutations characterized in this study found in the conserved regions of RNase H2 subunits are RNase H2A-G37S that corresponds to Rnh201-G42S in *S. cerevisiae*, and RNase H2C- R69W that corresponds to Rnh203-K46W in *S. cerevisiae*.

The mutations corresponding to human RNASEH2A-G37S and RNASEH2C-R69W in yeast are *rnh20*1- G42S and *rnh203*-K46W. To capture the rNMPs and the DNA upstream of the incorporated rNMPs we utilized the *ribose-seq* technique^22,23^, and performed the sequence analysis using the Ribose-Map bioinformatics toolkit^24–26^ to obtain the coordinates of rNMPs in the yeast nuclear and mitochondrial genomes. We determined embedded rNMP features like rNMP abundance rate, composition, and sequence preferences around the site of rNMP embedment in the DNA of cells carrying the AGS- orthologous mutations. Moreover, we compared the rNMP features of the AGS ortholog mutants with those found in the DNA of wild-type and *rnh201*△ mutant cells. We compared the presence of rNMPs nuclear vs. mitochondrial DNA to reveal changes in rNMP rates in the nucleus and thus sheds light on the catalytic activity of the RNase H2 enzyme complex. We also looked at the sequence context around the rNMP-incorporation sites in the yeast genome of the two AGS mutants, which provided insights on the effect of the AGS mutations on the functionality of the RNase H2-complex. Furthermore, we examined rNMP composition and sequence context of the most abundant rNMPs, as well as features of the rNMPs located around early firing ARS in the AGS ortholog mutants.

## RESULTS

### Increased rNMP-incorporation rates in the nuclear genome of *rnh201*-G42S mutant cells

With the goal to study rNMP-incorporation patterns in AGS mutants, we first engineered two AGS orthologous mutations occurring in conserved domains of human and yeast RNase H2 subunits. Two AGS mutations commonly found in human patients are RNASEH2A-G37S and RNASEH2C-R69W. The yeast ortholog mutants are *rnh201*-G42S and *rnh203*-K46W, respectively (Figure 1). RNASEH2A-G37S is in the catalytic site, whereas RNASEH2C-K69W is outside of the catalytic site but it is still very close to the Subunit A-catalytic site^10^. The yeast ortholog mutation *rnh201*-G42S and *rnh203*-K46W were created in *S. cerevisiae* in the BY4742 background (*MAT*α *his3*Δ1 *leu2*Δ0 *lys2*Δ0 *ura3*Δ0) and compared with the wild-type RNase H2 and *rnh201*△ genotypes. We built 14 ribose-seq libraries consisting of BY4742 wildtype, *rnh201*-G42S, *rnh203*-K46W, and *rnh201*△ genotypes. In addition to these libraries, in the analyses, we also included two wild-type and one *rnh201*△ ribose-seq libraries previously published derived from BY4742 backgrounds^16^. Multiple sets of restriction enzymes were used for in the preparation of different ribose-seq library, as shown in Table S<u>1</u>. The different combinations of restriction enzymes (RE) used to fragment the yeast genomic DNA were (i) RE1: DraI, EcoRV, and SspI; (ii) RE2: AluI, DraI, EcoRV, and SspI; and (iii) RE3: RsaI and HaeIII. The sequencing data of the ribose-seq libraries were analyzed using the Ribose-Map^24–26^ bioinformatics toolkit to first obtain the rNMP coordinates in the sacCer2 genome.

To study rNMP-incorporation rates in the yeast nuclear genome of each genotype, we looked at the percentage of rNMPs in the nuclear and mitochondrial DNA from the total rNMP count obtained for each rNMP library. We then compared these percentages in each library of all genotypes (wild-type, *rnh203*- K46W, *rnh201*-G42S, and *rnh201*△). Wild-type libraries have less than 15% rNMPs in the nucleus from the total genomic rNMPs (Figure 2A, Table S1). On the contrary, *rnh201*△ cells have > 70% rNMPs found in the nucleus. Libraries of the *rnh201*-G42S genotype show higher rNMP incorporation in the nucleus, like those of the *rnh201*△ genotype, suggesting impaired catalytic activity of RNase H2. Differently, the percentages of rNMPs found in the nuclear DNA of *rnh203*-K46W libraries are low, like those detected in the wild-type libraries suggesting little to no effect of the *rnh203*-K46W mutation on the catalytic activity of the RNase H2 complex.

**Figure 2.**
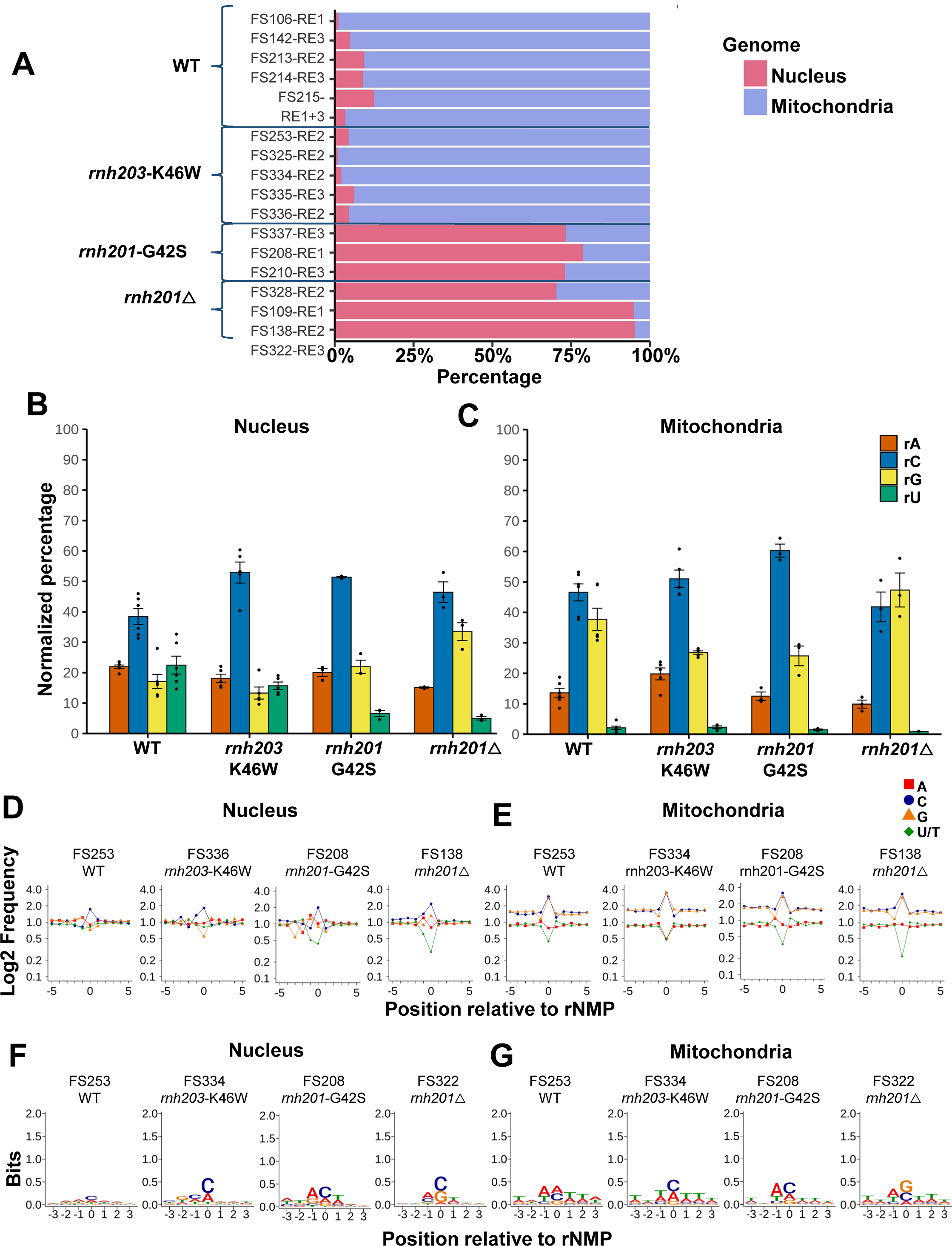
rNMP-incorporation rates and patterns in the nuclear and mitochondrial genome of *S. cerevisiae* AGS-ortholog mutants. (A) Horizontal stacked-bar plots showing rNMP-abundance rates as percentage of rNMPs found in the nuclear (red bars) and the mitochondrial (blue bars) genome of ribose-seq libraries (Table S1) of *S. cerevisiae* strain BY4742 with the indicated genotypes (wildtype, *rnh203*-K46W, *rnh201*-G42S, and *rnh201*△). (B) Bar graphs corresponding to the mean value and standard error of normalized percentages of rNMPs found in nuclear and mitochondrial (C) DNA of rNMP libraries for the indicated genotypes. The percentages are normalized to the base composition of the reference genome, such that the expected percentage for each rNMP base is 25 %^16^. Percentages without normalization of rNMPs are shown in Figure S1 and are listed in Table S2. (D) Sequence context +/- 5 nucleotides (nt) from the site of rNMP presence, which is indicated by the 0 position, for all rNMPs found in nuclear and (E) mitochondrial DNA, respectively. The frequency of each nucleotide is normalized to the frequency of the corresponding nucleotide present in the nuclear or mitochondrial reference genome. The plots shown are for one sample library of each genotype with the library and genotype indicated on top of the plots. The library name and genotype for the displayed data are indicated on top of each plot. Red square, A; blue circle, C; orange triangle, G; and green rhombus, U. The sequence plots generated from all rNMP libraries of this study are presented in Figure S2. (F) Sequence logo plots flanking +/- 3 nt from the rNMP position (0) of top 1 percentile locations with the highest rNMP counts observed in nuclear and (G) mitochondrial DNA. The *y*- axis shows the level of sequence conservation, represented in bits. The library name and genotype for the displayed data are indicated on top of each plot. The Sequence logo plots for all the libraries are present in Figure S3.

### Biased rC levels in *rnh201*-G42S and *rnh203*-K46W mutant genotypes

We normalized the rNMP-base composition with respect to the nucleotide base composition found in the reference genome. Bar plots of these normalized percentages of each rNMP base in the nucleus (Figure 2B) show higher rC percentages for *rnh201*-G42S and *rnh203*-K46W genotypes in comparison to the normalized percentage of rC in the wild-type genotype. The rNMP composition without normalization (Figure S1A) also indicates high rC percentages in AGS mutants amongst all other rNMP bases (rA, rG and rU). In nuclear DNA of *rnh201*-G42S libraries, alongside the increase in rC, there is decrease in rU like that seen in *rnh201*△ libraries (Figure 2B, Figure S1A). The biased rC level in nuclear DNA of *rnh201*- G42S and *rnh203*-K46W libraries is unique in comparison to the rNMP composition found in nuclear DNA of *rnh201*△ libraries, as *rnh201*△ libraries have biased rC and rG level in nuclear DNA. Normalized rNMP- base percentages in the mitochondrial genome (Figure 2C) for all genotypes have biased rC and rG normalized percentages as compared to rA and rU, consistent with results previously seen in yeast ribose-seq libraries^23^. rC does appear to be predominantly higher than rG in mitochondrial DNA of the *rnh201*- G42S and *rnh203*-K46W libraries as compared to wild-type and *rnh201*△ libraries (Figure 2C). rNMP composition without normalization in the mitochondrial genome shows rC amongst the most abundant ribonucleotide in *rnh201*-G42S and *rnh201*△ libraries unlike in wild-type and *rnh203*-K46W libraries. (Figure S1B).

### Specific sequence context upstream of rNMP incorporation sites in *S. cerevisiae* ortholog *rnh203*- K46W mutant libraries

To study the sequence context and nucleotide preference of rNMP incorporation site in every genotype, we calculated the deoxynucleotide frequencies around +/- 100 bases (zoomed out) and +/- 5 bases (zoomed in) near the unique sites of rNMP embedment for nuclear and mitochondrial genome in every library. After normalization to the mononucleotide frequencies observed in the reference genome, frequencies of deoxyribonucleotide monophosphates (dNMPs) around rNMP sites in the nuclear and mitochondrial genome show a preference (higher frequency) for dNMPs with base G or C in the mitochondria as compared to the nucleus where frequencies of A, C, G and T are almost similar (Figure 2D and E, and Figure S2A-D). This feature of rNMPs in the yeast mitochondrial genome is in line with what has been observed before with other budding yeast strains^16^ suggesting rNMP presence in GC clusters in the yeast mitochondrial genome^27^. Thus, despite the low percentage of dCMPs and dGMPs in mtDNA, rCs and rGs were often surrounded by dCMPs and dGMPs.

The sequence context near +/- 5 bases of the rNMP-embedment sites (0^th^ position) in nuclear DNA of the *rnh203*-K46W genotype shows preference for dCMP one base upstream (position -1) (Figure 2D). The dCMP preference at position -1 from the rNMP in *rnh203*-K46W libraries is particularly evident when the rNMP has base C or U (Figure S2A, B). Differently, in *rnh201*-G42S libraries, we see preference for dAMP one base upstream (position -1) of the rNMPs (Figure 2D), particularly when the rNMP has base A or G (Figure S2A, B). The rNMP frequency observed in the 0 position of the sequence plots is consistent with the normalized rNMP percentages for the respective genotypes (Figure 2B, C). Unlike for the nuclear genome, there is no observable difference in the frequencies of dNMPs around the rNMP site among the different genotypes for the mitochondrial genome. As previously observed for rNMPs in yeast mitochondrial DNA^16^, rCMP and rGMP sites are surrounded by dCMPs and dGMPs, respectively, within GC-rich regions (Figure 2E and Figure S2C, D).

### Unique rNMP-hotspot composition and consensus sequence in AGS-orthologous mutants

To analyze the sequence context around the highly frequent rNMP-incorporation sites in the yeast AGS- ortholog libraries, we obtained the hotspots of the rNMP counts present (1 percentile of highly abundant rNMP-incorporation sites) in nuclear and mitochondrial DNA. Using the Ribose-Map Hotspot Module, we obtained the genomic sequence of these abundant rNMP sites 3-nucleotides upstream and downstream of the rNMPs on the same strand as the rNMP, and visualized the results using the sequence logo plots (Figure 2F, G and Figure S3). We find that rCMP (0 position) is the dominant rNMP in the nucleus for all RNase H2-mutant genotypes tested (*rnh203*-K46W, *rnh201*-G42S and *rnh201*△), whereas there is no dominant rNMP seen in hotspots from wild-type nuclear libraries (Figure 2F). The rCMP hotspots are preferentially preceded by dCMP in *rnh203*-K46W libraries (-1 position), whereas in *rnh201*-G42S and *rnh201*△ libraries rCMP hotspots are preceded by dAMP and followed by dTMP (Figure 2F, Figure S3A). Sequence-logo plots of rNMPs in mitochondrial DNA do not reveal strong patterns in the AGS-mutant libraries as compared to wild-type or *rnh201*△ libraries, except that the rNMP position is less frequently an rGMP (Figure 2G, Figure S3B). Hotspots of rNMPs in mitochondrial DNA of all genotypes studied show an evident preference for dAMP at position -1 and a fit within the consensus TNArSWTW sequence previously identified^16^ (Figure 2G, Figure S3B).

We also looked at hotspots shared within at least 2 libraries of each genotype and selected the most abundant rNMP-incorporation sites (top 75 in nucleus and top 25 in mitochondria) using the rNMP- enrichment factor of rNMP counts per base relative to genomic rNMPs per base (see Methods). The counts used were normalized based on the relative base coverage obtained from DNA sequenced for each library to exclude any false positive hotspots in low complexity (repeat) regions of the yeast genome. The top-75 rNMP sites of the shared hotspots in the nucleus and the top-25 rNMP sites of the shared hotspots in mitochondria are mapped with their base (Figure 3). The +/- 3-bp flanking sequence consensus and the composition of hotspots normalized to the nucleotide frequency in the nuclear and mitochondrial genome were also studied (Figure S4A, B) with overlapping gene annotations (Table S4 and S5). The shared hotspots in nuclear DNA of RNase H2 wild-type cells do not appear to cluster around a specific location, whereas we detect more than 15 rNMP-hotspot sites in a cluster on Chr IV between 0.5 Mb to 1 Mb in *rnh203*-K46W libraries. Interestingly, the top-75 hotspots in nuclear genome of *rnh203*-K46W have higher rC than wildtype (Figure 4A, B). The top-75 hotspots in the nuclear genome of *rnh201*-G42S mutation have more than 50 % of hotspot sites with rC, and *rnh201*△ libraries have the highest percent of rG hotspots (Figure 3 C, D, Figure S4C, and Table S4E). We also see that *rnh203*-K46W libraries have a similar level of rA as wild-type libraries, but a lower level of rG like *rnh201*-G42S libraries, making the hotspot composition in both AGS mutants unique in comparison with the wild-type and *rnh201*△ libraries (Figure 3D, Figure S4A, C, and Table S4E). This observation is also validated using sequence logo plots for the top 2 and 5 percentile highly abundant rNMP locations from all unique locations in nuclear and mitochondrial DNA for each genotype studied (Figure S4A, B). In nuclear DNA, we do not detect a strong consensus in the rNMP position (0 position) in wild-type libraries. Instead, the rC is predominant in all mutant genotypes, and rG is more abundant in *rnh201*△ (Figure S4A). We have also mapped the genes and their types to top 75 shared common hotspots in all genotypes to find that more than 69 hotspots out of 75 hotspots map to protein coding genes in *rnh201*-G42S where as in *rnh203*-K46W only 51 hotspots map to protein coding genes (Table S4B, C). Increase in hotspots mapping to protein coding genes is also seen in *rnh201*△ (58 hotspots, Table S4D) vs wild type (49 hotspots, Table S4D). The top-25 rNMP sites of the shared hotspots in mitochondria show that the rC site is more abundant in *rnh203*-K46W and *rnh201*-G42S libraries compared to wild-type and *rnh201*△ libraries (Figure 3E-H and Table S5E). By analyzing the consensus sequence of the top-2 and 5 percentiles of rNMP hotspots, we detect more rA and rC in all genotypes with rC being higher in rnh201-G42S, and rG being abundant only in *rnh201*△ (Figure S4B).

**Figure 3.**
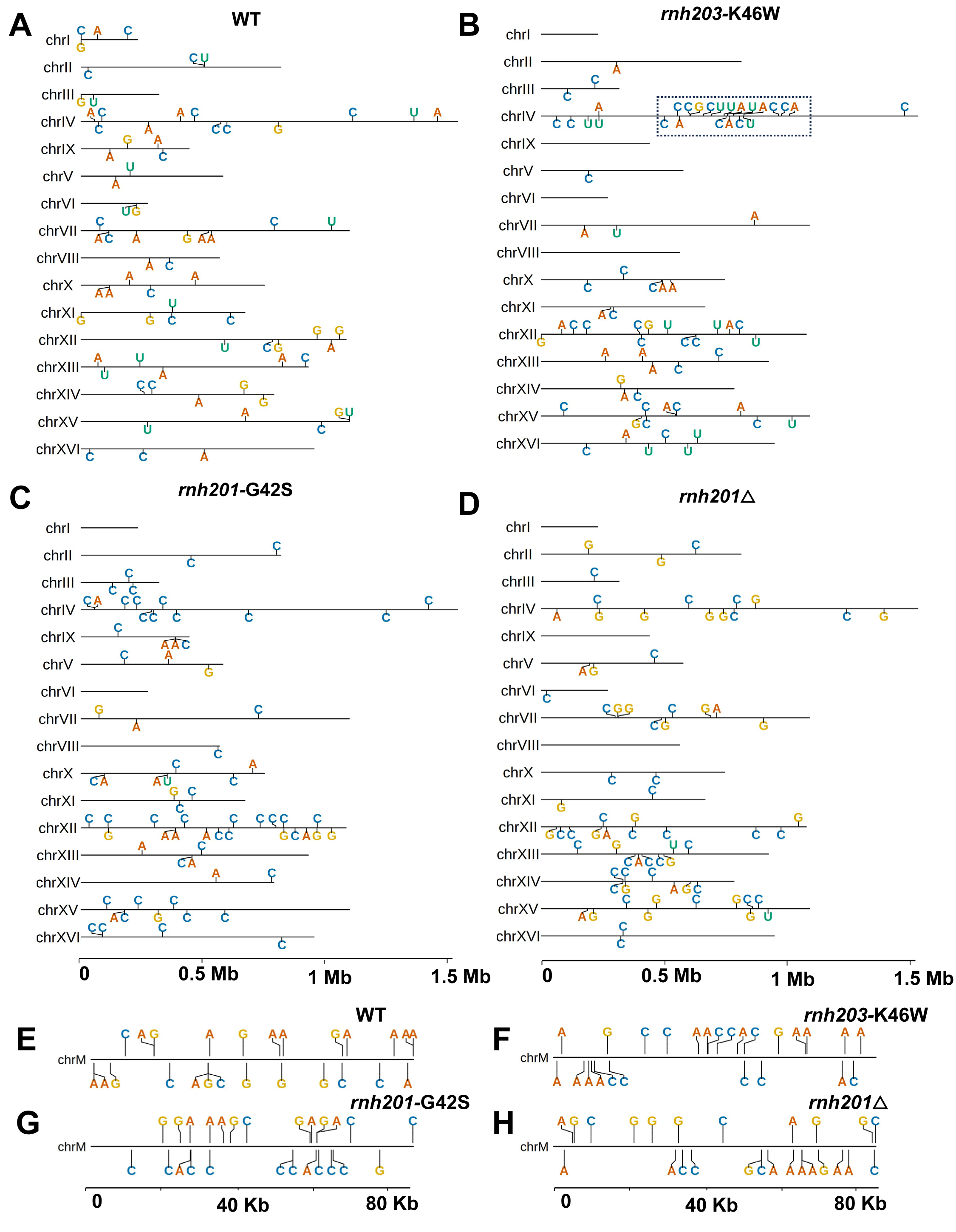
Most abundant shared hotspots in nuclear and mitochondrial DNA of the *rnh203*-K46W and *rnh201*-G42S AGS ortholog mutants Genome-mapped locations of the 75 most abundant shared rNMP hotspots in nuclear DNA of ribose-seq libraries of (A) wild-type, (B) *rnh203*-K46W, (C) *rnh201*-G42S, and (D) *rnh201*△ genotype. Genome mapped locations of the 25 most abundant shared rNMP hotspots in mitochondrial DNA of ribose-seq libraries of (E) wildtype, (F) *rnh203*-K46W, (G) *rnh201*-G42S, and (H) *rnh201*△ genotype. The represented annotations for rA, rC, rG, and rU are shown in red, blue, yellow, and green color letters, respectively. The shared hotspots are selected based on occurrence in at least 2 libraries in each genotype and the highest rNMP Enrichment Factor (ratio of rNMPs to average rNMP per base in the respective library genome). Chromosome locations +/- 3 nt of flanking sequence, enrichment factor, mapped genes, and composition of shared hotspots is provided in Table S4 for the shared hotspots in the nuclear genome and in Table S5 for the shared hotspots in the mitochondrial genome. Shared hotspot clusters identified on Chr IV in *rnh203*-K46W libraries are indicated by a dashed frame. Sequence logo plots for top 75 (nucleus) and top 25 (mitochondria) with top 2 and 5 percentile hotspots of unique rNMP locations (both nucleus and mitochondria) in each genotype are shown in Figure S4.

**Figure. 4.**
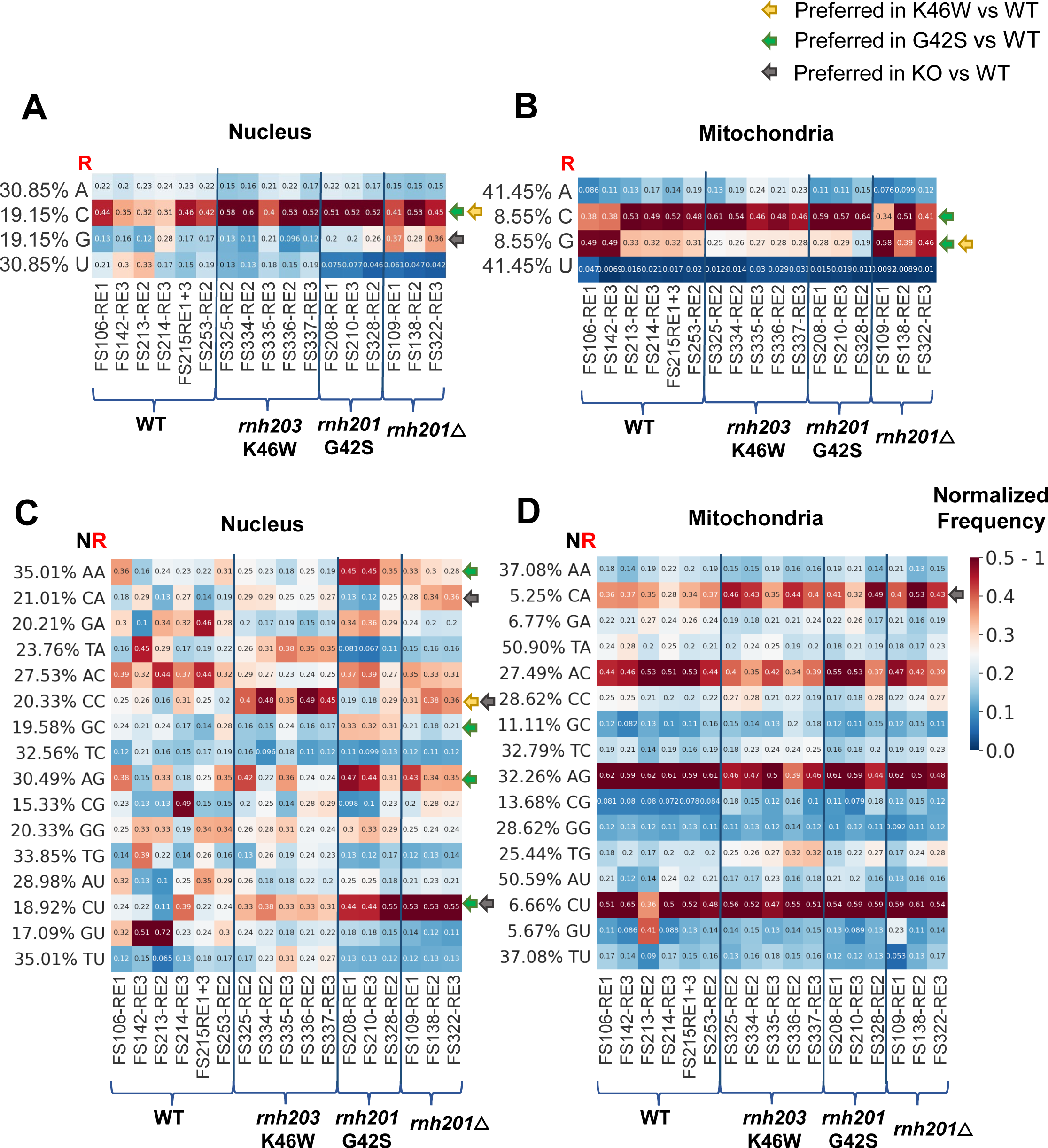
rNMP base composition and dinucleotide patterns identified in the *rnh203*-K46W and *rnh201*-G42S AGS orthologs mutants. Heatmap of normalized frequency of each type of rNMP (rA, rC, rG, and rU) in nuclear (A) and mitochondrial (B) DNA for all the ribose-seq libraries of this study. Heatmap analyses with normalized frequency of (C) nuclear and (D) mitochondrial NR dinucleotides (rA, rC, rG, and rU with the upstream deoxyribonucleotide with base A, C, G, or T) for all the ribose-seq libraries of this study. The formulas used to calculate these normalized frequencies are shown and explained in Methods. Each column of the heatmap shows results of a specific ribose-seq library. Each library name is indicated underneath each column of the heatmap with its corresponding restriction enzyme (RE) set used. The ribose-seq libraries of the same genotype are also grouped together by brackets and separated by thick vertical blue lines. Each row shows results obtained for rNMP-mononucleotide or dinucleotide combination. The actual percentages of rNMP (rA, rC, rG, and rU) or dinucleotides of fixed base A, C, G, or T for the indicated base combinations (AA, CA, GA, and TA; AC, CC, GC, and TC; AG, CG, GG, and TG; and AT, CT, GT, and TT) present in the nuclear and mitochondrial genome of *S. cerevisiae* are shown to the left of the corresponding heatmaps. The observed % of rNMPs or dinucleotides with rNMPs with base A, C, G, or U were divided by the actual % of each rNMP or dinucleotide with fixed base A, C, G, or T in nuclear or mitochondrial DNA. The bar to the right shows how different frequency values are represented as different colors: white for 0.25; light red to red for 0.25 to 0.5–1, and dark blue to light blue for 0.25 to 0. Yellow, green, and grey arrows indicate the nucleotide frequency significantly preferred for *rnh203*-K46W, *rnh201*- G42S, and *rnh201*△ mutant libraries, respectively, in comparison to wild-type libraries. Significantly different frequencies are highlighted based on one tailed Mann Whitney-Wilcoxon *U* test between nucleotide frequencies in the AGS mutant genotypes *vs*. the expected value of 0.25 (Table S6 and 7), as well as two tailed Mann-Whitney Wilcoxon *U* test AGS-mutant libraries *vs*. wild-type libraries (Table S8 and 9). Dinucleotide heatmaps for the most frequent dNMP found downstream of the rNMPs in the nuclear and mitochondrial DNA are presented in Figure S5.

### Sequence-specific preferences in *S. cerevisiae* orthologs of RNase H2C-R69W and RNase H2A- G37S mutants

To further investigate and validate the sequence-context conservation of the rNMPs and their upstream and downstream dNMPs in the *S. cerevisiae* orthologs of human RNase H2C-R69W and RNase H2A-G37S mutants, we obtained heatmaps of the normalized frequencies of each rNMP mono- and di- nucleotide combination in nuclear and mitochondrial DNA (Figure 4A-D). We used one-tailed Mann-Whitney *U* test to determine the significant preference of each combination (Table S6 and S7) and two tailed Mann-Whitney *U* test to determine significantly different frequencies between any of the RNase H2- mutant libraries and the wild-type libraries (Table S7 and S8). Significant differences in the pairwise comparisons with *P* value < 0.05 are highlighted with yellow arrows for *rnh203*-K46W libraries, green arrows for *rnh201*-G42S libraries and grey arrows for *rnh201*△ libraries. The mononucleotide-heatmap analysis of rNMPs in nuclear DNA shows that the normalized frequency of rC increases significantly (*P* value < 0.05 in Table S6) in both AGS-mutant genotypes, *rnh203*-K46W and *rnh201*-G42S (Figure 4A); whereas, in the mitochondrial DNA, rC increases in *rnh201*-G42S as compared to wild-type and *rnh201*△ libraries (Figure 4B, Table S7). Moreover, in mitochondrial DNA, while rG is still preferred in comparison to rA and rU in both the AGS-mutant libraries (*P* value < 0.05 in Table S7), the frequency of rG decreases in both *rnh201*-G42S and *rnh203*-K46W as compared to wild-type and *rnh201*△ libraries, (Figure 4B, *P* value < 0.05 in Table S9).

In the dinucleotide-heatmap (NR) analysis with the rNMP (R) and its upstream dNMP (N), NR combinations in the nucleus, CrC has the strongest frequency in *rnh203*-K46W as compared to libraries of the other genotypes, in which we observe preference for ArC (Figure 4C). For the *rnh201*-G42S libraries, we see NR-dinucleotide combinations with preference for dAMP: ArA, ArC, and ArG, and upstream for dGMP: GrA, GrC, and GrG (Figure 4C and *P* value < 0.05 in Table S6). Among these dinucleotide preferences in the *rnh201*-G42S libraries ArA, ArG, and GrC frequencies are significantly different (*P* value < 0.05 in Table S8) from those of the wild-type and *rnh201*△ libraries (Figure 4C and Table S8). We also detect a strong CrU preference in *rnh201*-G42S libraries as in *rnh201*△ libraries. The patterns for ArG and CrU in *rnh201*-G42S are consistent with the patterns observed in *rnh201*△ libraries, whereas *rnh201*-G42S libraries have a higher preference for GrA, GrC, and GrG and a lower preference for CrA, CrC, and CrG compared to *rnh201*△ libraries (Figure 4C, *P* value < 0.05 in Table S6). Dinucleotide preferences in the mitochondrial genome (Figure 4D) are very similar for all genotypes, with a markedly conserved preference for CrA, ArC, ArG, and CrU, partially like those observed in nuclear DNA of *rnh201*△ libraries (Figure 4C, D, *P* value < 0.05 in Table S7). However, there are some differences observed in the frequencies between genotypes, i.e., lower frequency of ArC and ArG in *rnh203*-K46W libraries as compared to wild-type libraries (Figure 4D, *P* value < 0.05 in Table S9).

In the heatmap analysis of the rNMP with its downstream dNMP, dinucleotide (RN), we detect weak preferences in both nuclear and mitochondrial DNA (highlighted using arrows in Figure S5 and *P* value < 0.05 in Table S8 and S9). The dinucleotide rUG is preferred in nuclear DNA of *rnh203*-K46W libraries being significantly higher than in wild-type libraries (highlighted using yellow arrows in Figure S5, *P* value < 0.05 in Table S6 and 8). The dinucleotides rAT, rCA, and rGT are preferred in the nuclear DNA of *rnh201*-G42S libraries compared to wild-type libraries (highlighted using green arrows in Figure S5, *P* value < 0.05 in Table S6 and 8). The frequencies of dinucleotides rAT and rGT frequencies in *rnh201*- G42S libraries are like those in *rnh201*△ libraries (highlighted using grey arrows in Figure S5, *P* value < 0.05 in Table S6 and S8).

### *S. cerevisiae* orthologs of RNase H2C-R69W and RNase H2A-G37S mutants show specific dinucleotide patterns on the leading and lagging strand around ARS sequences of the yeast genome

To examine whether the yeast AGS-ortholog mutant libraries display similar or different strand bias patterns on the leading and lagging strands, we analyzed rNMP distribution and patterns on the leading and lagging strands within windows of 0-4 kb and 4-10 kb from ARSs in the genome of wild-type, *rnh203*-K46W, *rnh201*-G42S, and *rnh201*△ strains of this study (Figure 5). We found that as in wild-type libraries^17^, the *rnh203*-K46W libraries tend to have a slight increased level of rNMPs on the leading strand. Such preference for the leading strand, as in *rnh201*△^17^, is more evident in *rnh201*-G42S libraries in both 0-4 kb and 4-10 kb flanking regions of the ARSs (Figure 5A, B). In the mononucleotide-heatmap analysis of rNMPs present on the leading vs lagging strands in the 4-10-kb region around early-firing ARSs, we do not detect strand-biased composition for the AGS mutant libraries, as detected for wild-type and *rnh201*△ libraries of wild-type mutant Pols^17^. It has been found that rNMPs have markedly distinct dinucleotide frequencies on the leading and lagging strands 4-10-kb upstream and downstream of early-firing ARSs in rNMP libraries of *rnh201*△ cells expressing wild-type or mutant forms of DNA polymerase (Pol) δ or Pol ε. Leading and lagging strand-dinucleotide patterns show signatures of incorporation due to the activity of DNA Pol ε and Pol δ on these strands, respectively. The normalized dinucleotide frequencies of ArA, ArC, and ArG are higher on the leading strand, while those for CrA, CrC, and CrG are higher on the lagging strand^17^ (see grey frames for *rnh201*△ in Figure 5E, F). In the heatmap analysis of dinucleotide preferences in nuclear DNA, we found that for *rnh203*-K46W libraries the preferred dinucleotide CrC has higher frequency on the lagging strand as compared to leading strand in the 4-10 kb region from early ARSs (Figure 5E, F; see yellow frames). This pattern of CrC being higher on the lagging vs the leading strand, likely due to rNMP incorporation by Pol δ activity, is also seen in *rnh201*△ libraries, but it is barely present in *rnh201*-G42S libraries (Figure 5E, F). For *rnh201*-G42S libraries, we detect preference for ArA, ArC, and ArG dinucleotide patterns in 4-10-kb region of the leading strand (yellow frames) as compared to the lagging strand, as also detected in the *rnh201*△ libraries, showing incorporation patterns due to Pol ε activity (Figure 5E, F).

**Figure 5.**
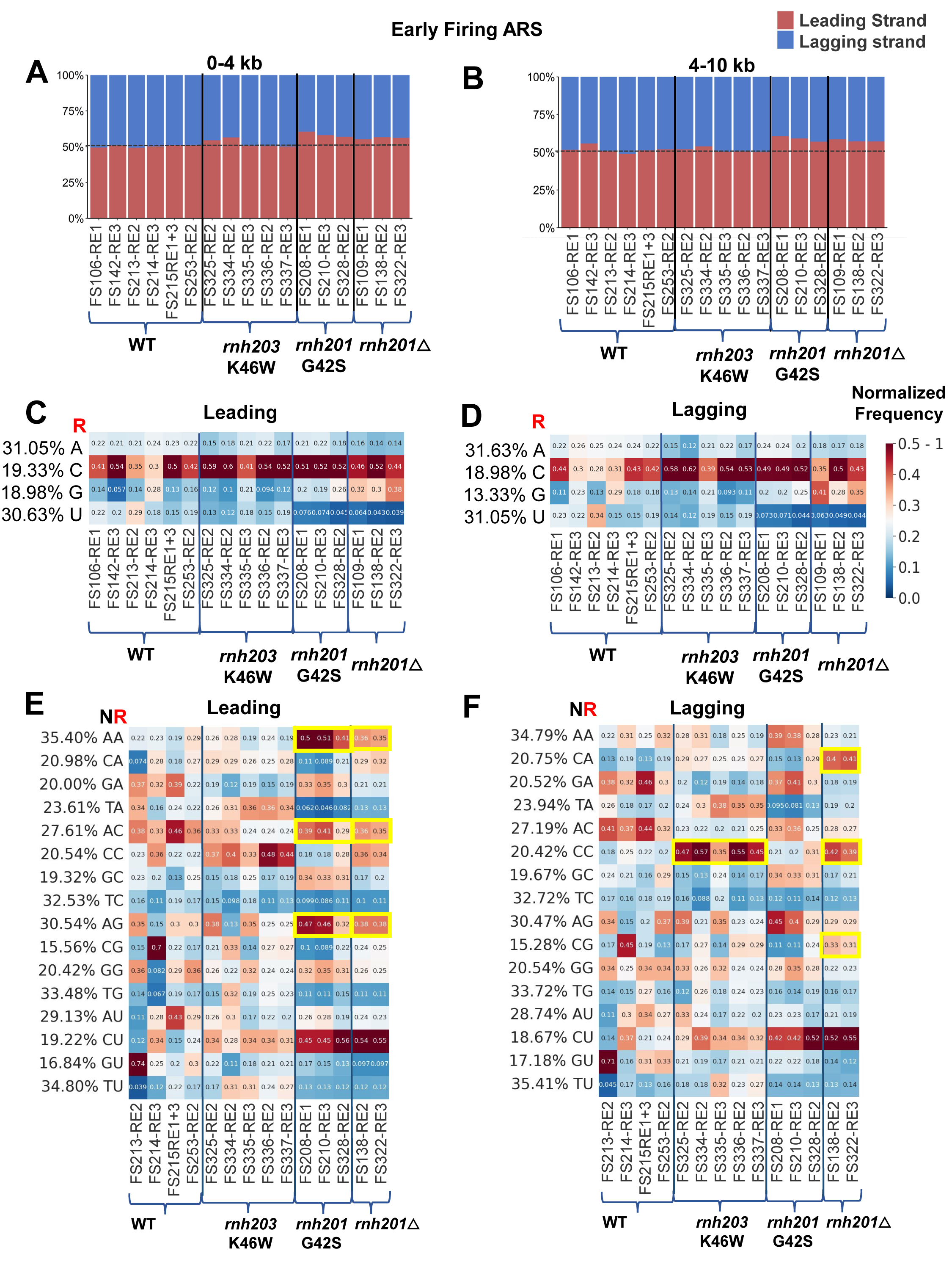
The *rnh203*-K46W and *rnh201*-G42S AGS orthologs mutants display bias distribution, composition, and NR-dinucleotide patterns on the leading and lagging strand in yeast nuclear DNA. Stacked-percentage bar plots of rNMP counts in (A) 0 – 4,000 nt and (B) 4,000 – 10,000 nt windows on the leading (red bars) and lagging (blue bars) strand regions adjacent to early firing ARSs. Mono nucleotide heatmaps analysis in 4,000 – 10,000 nt windows in the leading (C) and lagging (D) strands with frequencies normalized to the dNMP content in the 4000 – 10,000-nt window around the early-firing ARSs on the leading or lagging strand, respectively. Dinucleotide NR (rA, rC, rG, and rU with the upstream deoxyribonucleotide with base A, C, G, or T) heatmap analysis to reveal preferences on the (E) leading and (F) lagging strand for wild-type, *rnh201*△, and AGS mutants 4,000 -10,000 nt from the early-firing ARSs. The formulas used to calculate these normalized frequencies are shown and explained in the Methods. Each column of the heatmap shows results of a specific ribose-seq library. Libraries with more than 400 rNMPs in the leading and lagging strand are displayed in the heatmaps. Each library name is indicated underneath each column of the heatmap with its corresponding restriction enzyme (RE) set used. The ribose-seq libraries of the same genotype are also grouped together by brackets and separated by thick vertical blue lines. Each row shows results obtained for rNMP or dinucleotide combination. The actual percentages of rNMP (rA, rC, rG, and rU) dinucleotides of fixed base A, C, G, or T for the indicated base combinations (AA, CA, GA, and TA; AC, CC, GC, and TC; AG, CG, GG, and TG; and AT, CT, GT, and TT) present in the 4,000 – 10,000-nt windows around early-firing ARSs on the leading and lagging strands are shown to the left of the heatmaps. The observed % of rNMPs and dinucleotides with rNMPs with base A, C, G, or U were divided by the actual % of each rNMP and dinucleotide with fixed base A, C, G, or T in 4,000 – 10,000-nt windows around ARSs on the leading and lagging strands. The bar to the right shows how different frequency values is represented as different colors: white for 0.25; light red to red for 0.25 to 0.5–1, and dark blue to light blue for 0.25 to 0. Yellow boxes indicate preferences with higher frequencies found in ArA, ArC and ArG in the leading strand, and CrA, CrC, and CrG found in the lagging strand for AGS mutants *rnh203*-K46W, *rnh201*-G42S, and *rnh201*△.

## DISCUSSION

### Rnh201*-*G42S (RNase H2A-G37S ortholog) but not Rnh203-K46W (RNase H2C-R69W ortholog) has reduced RER activity

The G37S mutation in human RNase H2A has been identified in individuals with AGS and is in the highly conserved catalytic site of the RNase H2 enzyme^28^.This specific mutation affects the catalytic activity of the RNase H2 enzyme, impairing its ability to cleave rNMPs^29^ and process RNA/DNA hybrids effectively^30^. In our analyses, the RNase H2A-G37S mutant ortholog in yeast, Rnh201-G42S, reflects lack of catalytic activity because of the highly increased rNMP percentage in nuclear DNA relative to the rNMP presence in mitochondrial DNA, as observed for *rnh201*△ libraries (Figure 2A). The R69W mutation in the human RNase H2C subunit affects the RNase H2-complex stability^10,31^, and it is located very close to the catalytic site of RNase H2A within the structure of the RNase H2 complex. The yeast ortholog of RNase H2C- R69W is Rnh203-K46W. The ribose-seq libraries of *rnh203*-K46W, unlike those of *rnh201*-G42S and *rnh201*△ cells, have less than 10 % of rNMPs in the nuclear genome with most rNMPs found in the mitochondrial genome like wild-type libraries. Because RNase H2 is not active on mitochondrial DNA^7,32^ these results indicate that the RNase H2 catalytic activity is mostly intact in the *rnh203*-K46W but compromised in the *rnh201*-G42S genotype. There are numerous studies that infer rNMP abundance in the genome of cells by sensitivity test for stress-inducing agents^29,33^ or nick translation reaction^34^ that provide information of rNMP levels. By using the ribose-seq technique to directly capture rNMP sites in genomic DNA and paired analysis with the Ribose-Map toolkit to map them at single-nucleotide resolution, our study specifically enables to obtain specific location, composition of rNMPs, distribution, and sequence context patterns of rNMP sites in yeast AGS-ortholog mutants, which can help to understand the consequences of the orthologous mutations in humans. In future studies it will be possible to apply the same techniques to directly capture and study rNMPs embedded in the DNA of AGS-mutant samples derived from cell lines or patients.

### Higher rC and rG frequency content in the nuclear genome of the rnh201-G42S and the rnh203- K46W mutant cells

Previous research investigating the composition of rNMPs in yeast DNA reported high rC frequency in the nuclear genome for all strains studied, including the BY4742 background, in wild-type and *rnh201*△ cells ^16^. We found consistent trends in our study with even higher level of rC in the nuclear genome of both AGS-ortholog mutants (Figure 2C). In line with results in yeast mitochondrial DNA, in which rG tends also to have a high frequency, particularly in the BY4742 strain^16^, *rnh201*△ cells display a higher frequency of rG in nuclear DNA^16^ (Figures 2B and 4A). Between the two AGS-ortholog mutants studied here, *rnh201*- G42S mutant has nuclear rG frequency higher than wild-type cells, unlike *rnh203*-K46W, like that observed in *rnh201*△ cells. This result, in accordance with the higher abundance of nuclear rNMPs in *rnh201*-G42S cells compared to *rnh203*-K46W cells (see Figure 2A) discussed above, provides further evidence that the cleavage activity of RNase H2 is compromised in the *rnh201*-G42S but not in the *rnh203*-K46W cells. Another composition pattern observed for the lack of RNase H2 activity on rNMPs is the low level of rU in nuclear DNA of *rnh201*-G42S libraries as compared to wild-type libraries and almost as low as in the *rnh201*△ libraries (Figure 2B and 4A). Even lower frequency of rU is observed in the mitochondrial genome (Figure 2C and 4B), in which there is no RNase H2 activity^18,19^. However, the level of nuclear rU in the *rnh203*-K46W libraries is not different from that observed in wild-type libraries. Overall, these results suggest that the RNase H2-complex activity is mainly retained in the *rnh203*-K46W mutant cells, and while not completely abolished, it is severely compromised in the *rnh201*-G42S cells allowing abundant rNMP presence in the nuclear DNA.

### rNMP hotspots in the yeast AGS-ortholog mutants show unique composition and distribution as compared to wild-type and rnh201△ samples

Hotspots, top one percentile locations in the genome with highest rNMP counts in each library (Figure 2C, D and Figure S3), and shared hotspots, rNMP locations with highest average rNMP enrichment recurring in at least two libraries of same genotypes (Figure 3 and Figure S4), show trends in composition mostly consistent with those of all genomic rNMPs in the different genotypes. Nuclear rNMP hotspots found in wild-type libraries do not show preference for any rNMP (Figure 2C); whereas *rnh201*-G42S and *rnh203*- K46W libraries have a high level of rC hotspots and *rnh201*△ libraries have rC and rG hotspots. A high level of rC in the top 75 shared hotspots is found in the AGS mutants like in *rnh201*△ (Figure 3A-D and Figure S4C). Moreover, in *rnh203*-K46W libraries, the shared hotspots with rU are more abundant and those with rG and are much more prevalent than in *rnh201*△ and in *rnh201*-G42S libraries, as in wild-type cells (Figure 3A-D, Figure S4C). Both AGS-orthologous mutants have a low-level pf rG among the hotspots (Figure 3A-D, Figure S4C) and have a unique composition profile. The AGS-ortholog mutants have higher rC shared hotspots than wild-type and *rnh201*△ cells in both nuclear and mitochondrial DNA (Figure S4, Figure S4C, D). The *rnh203*-K46W cells have higher rA and lower rG hotspots than the *rnh201*- G42S cells in both nuclear and mitochondrial DNA (Figure S4C, D). Furthermore, the distribution of *rnh203*-K46W shared hotspots is clustered on chromosome IV within the 500,474 and 1009,648 loci (Figure 3B, Table S4). The different rNMP composition and distribution of the shared hotspot in the different RNase H2-genotype libraries compared to those detected in the wild-type RNase H2 libraries reveal rNMP sites that cannot be processed by the mutant forms of RNase H2 allowing their accumulation. Defective RNase H2A genotypes (*rnh201*-G42S and *rnh201*△) have more shared hotspots mapping to protein coding regions in the nucleus as compared to wild-type cells (Table S4).

### Both yeast AGS-ortholog mutants *rnh201*-G42S and the *rnh203*-K46W show distinct rNMP patterns suggesting distinct activity on rNMPs embedded in yeast nuclear DNA

The dinucleotide heatmap analysis of the nuclear rNMPs and their upstream dNMP clearly show distinct sequence patterns in each of the two types of AGS mutant libraries relative to wild-type and *rnh201*△ libraries, differently from the conserved common rNMP patterns obtained from analysis of mitochondrial DNA in all genotypes (Figure 4C and D). The *rnh20*3-K46W libraries show preference for dCMP upstream of rC (CrC), while *rnh201*-G42S libraries show preference for ArA, GrC, ArG, and CrU dinucleotide patterns. The difference between dinucleotide patterns in nuclear DNA of the two AGS mutants suggest that the mutations *rnh201*-G42S and *rnh20*3-K46W can change not just the activity of the RNase H2 enzyme but potentially also the binding position of the RNase H2 enzyme in nuclear DNA. This observation is also supported by the fact that the ArA pattern is more frequent in the *rnh201*-G42S libraries than in the *rnh201*△ libraries. Previous work that investigated the thermal complex stability^11^ of different RNase H2 mutants associated with AGS showed a low repair activity for the RNase H2A-G37S mutant, and a low complex stability for both the RNase H2A-G37S and the RNase H2C-R69W mutants.

While we find strong dNMP preferences upstream of the nuclear rNMP sites, the dNMP preference downstream of the nuclear rNMP sites are less conserved (Figure S5). However, we detected an apparent preference for the dGMP following rA and rU in libraries of *rnh203*-K46W cells. Such dinucleotide- downstream patterns may reflect an altered DNA binding capacity of the RNase H2 complex, potentially biased to specific sequence contexts. The dinucleotide patterns not only reflect the type of rNMPs that are not removed by RNase H2, but also the type of rNMPs that are incorporated by DNA polymerases. Thus, we also investigated the dinucleotide patterns around early firing ARSs in the budding yeast genome of the AGS-ortholog mutants, wild-type, and *rnh201*△ libraries. We studied the rNMP-dinucleotide patterns separately on the leading and the lagging strands in 4-10 kb-window regions flanking known early firing yeast ARSs as described previously^17^. The analyses confirmed a leading strand preference for dAMP upstream of rA, rC, rG, and in part rU, and a lagging stand preference for dCMP upstream of any rNMP in *rnh201*△ cells, as previously found^17^. Since a DNA Pol ε mutant that incorporates more rNMPs shows bias for rNMPs preceded by dAMP on the leading strand, and vice versa, a DNA Pol δ mutant that incorporates more rNMPs shows bias for rNMPs preceded by dCMP on the lagging strand, the ArN- dinucleotide pattern mainly reflects Pol ε-incorporated rNMPs, and the CrN-dinucleotide pattern mainly reflects Pol δ-incorporated rNMPs^17^. Interestingly, the dinucleotide pattern CrC not only is stronger in the *rnh203*-K46W libraries than in any other genotype libraries (Figure 4C), but it is also accentuated on the lagging compared to the leading strand of the *rnh203*-K46W libraries (Figure 5C, D), suggesting that the Rnh203-K46W mutant of RNase H2 does not efficiently process rNMPs introduced by Pol δ, especially rC, but it does process rNMPs introduced by Pol ε. On the contrary, the *rnh201*-G42S libraries show a marked preference for dAMP upstream of the rNMPs that is markedly accentuated on the leading strand, indicating that the *rnh201*-G42S mutant of RNase H2 may not be able to cleave effectively leading-strand rNMPs where DNA Pol ε is dominant.

### RNase H2 mutations in other diseases and cancer

Understanding the role of RNase H2 and the consequences of rNMP incorporation in DNA has broader implications beyond AGS. Characterizing mutations in yeast AGS ortholog mutants of RNase H2 aids in enhancing our comprehension not just of AGS mechanisms but also of genome maintenance processes and the repercussions of DNA repair deficiencies on human health. Our study also reflects how RNA processes, DNA repair and DNA replication are connected, offering insights into mechanisms of other diseases associated with dysregulations like those of AGS. RNase H2 impairment and mutations are observed in other conditions like systemic lupus erythematosus (SLE), also an autoimmune disorder characterized by the presence of nucleic acid and aberrant type I IFN signature^35,36^, and Werner syndrome (progeria), an inherited disorder marked by rapid aging and increased risk of cancer^37^.

It is interesting to note that the defects in RNaseH2A and H2B subunits are reported to cause inflammatory response^20,38^, whereas RNase H2 defects coupled with TP53 mutations lead to additional mutations, making it even more relevant to study roles of RNaseH2 in carcinogenesis^38^. In a recent study, RNaseH2C has also been reported to be a novel metastasis susceptibility factor in breast cancer^39^. On the contrary, RNaseH2A has also been shown to be important for sustaining cancer proliferation^40–45^ and knockout of RNase H2 can have tumor suppressive effects^38,46^. By studying embedded rNMP patterns, we may be able to decipher the effectiveness of RNaseH2A enzyme and rNMP removal in different types of cells. In the way RNASEH2A gene has been deemed to act as a prognostic Biomarker in many cancers^42^, rNMP features also can play a predictive role to suggest the state of cells and progression of several diseases including cancer, AGS, and other immune related diseases. Considering the multifaceted roles of the RNase H2 subunits in preventing human diseases, it is becoming more and more important to study effects of different mutations in the RNase H2-subunit genes and how rNMP incorporation is related to human disease to open new possibilities for disease prevention, early-stage detection or disease interventions. This study has not only helped identify embedded rNMP features in faulty RNase H2 but has also provided guidance for future research on rNMP features in the broader human genome across various cell types.

## Limitation

Here, we have presented yeast models of two AGS mutations in RNaseH2 in a pioneer work to understand the different rNMP characteristics found in the genomic DNA of two ortholog mutants of human RNaseH2, RNaseH2A-G37S and RNaseH2C-R69W. The analyses were performed in yeast gene orthologs *rnh201*- G42S and *rnh203*-K46W corresponding to the AGS mutants RNaseH2A-G37S and RNaseH2C-R69W in human RNaseH2, respectively. It would be interesting to explore patterns and hotspots of rNMPs embedded in the DNA of human cell lines or cells derived from patients carrying the same AGS mutations, as well as other AGS mutations. Complementary studies in human cells carrying AGS mutations would help to have a comprehensive understanding of the functions and consequences of the rNMPs embedded in the genome and their impact on DNA and RNA metabolism of the mutant cells. Due to the rare cases of AGS patients and lack of knowledge on embedded rNMP features in human nuclear genome, yeast was used as the eukaryotic model to determine the feasibility of expanding the work to human genome, and it is possible that not all features in AGS-orthologous mutants in yeast can be extrapolated to AGS mutants in humans.

## ACKNOWLEDGEMENTS

We thank S. Biliya, N. Djeddar, and A. Bryksin from the Molecular Evolution Core for advice and support with high-throughput sequencing, and the Partnership for an Advanced Computing Environment (PACE) at the Georgia Institute of Technology for their research cyberinfrastructure resources and services.

We acknowledge S. Randhawa, Y. Lee and M. Sun for critically reading the manuscript; and all members of the Storici laboratory for assistance and feedback on this study.

We acknowledge funding from the National Institute of Health, the National Institute of Environmental Health Sciences (R01ES026243 to F.S.), the Howard Hughes Medical Institute Faculty Scholar grant 55108574 (F.S.), the Mathers Foundation (AWD-002589 to F.S.), and the W. M. Keck Foundation (to F.S.). to support this work.

## AUTHORS CONTRIBUTIONS

Conceptualization: Francesca Storici, Deepali L. Kundnani, Taehwan Yang, and with help of Alli L. Gombolay

Methodology: Deepali L. Kundnani, Taehwan Yang with help from Alli L. Gombolay, Kuntal Mukherjee, Gary Newnam, Chance Meers, Zeel Mehta, Celine Mouawad and Francesca Storici

Investigation: Deepali L. Kundnani, Alli L. Gombolay, Taehwan Yang and Francesca Storici Data Curation: Deepali L. Kundnani with help of Taehwan Yang and Alli L. Gombolay Visualization, Formal Analysis and Validation: Deepali L. Kundnani

Interpretation: Deepali L. Kundnani and Francesca Storici Writing - Original Draft: Deepali L. Kundnani

Writing – Review & Editing: Deepali L. Kundnani, Taehwan Yang and Francesca Storici Funding Acquisition, Resources and Supervision: Francesca Storici

## TABLE TITLES AND LEGENDS

Table S1. AGS library information

Background, Strain, genotype, Library ID, fragmentation enzymes used, Library barcode, source of library, rNMP counts and percentages in nucleus and mitochondria of each library with mean and std for each genotype used in this study are mentioned.

Table S2. Oligonucleotides used in this study.

Name, length, and sequence of oligonucleotides used in this study are presented. All bold letters in the PCR primers indicate the specific sequence of index used in sequencing. P and Am indicate end modifications of phosphate and amino modifier, respectively. All oligonucleotides were desalted, except those marked with an asterisk (*), which were HPLC purified. All oligonucleotides were synthesized by Integrated DNA Technologies.

Table S3. AGS library rNMP composition in Nucleus and Mitochondria

Background, Strain, genotype, Library ID and composition of each rNMP base (rA, rC, rG and rU) are provided with mean and std for each rNMP base in each genotype.

Table S4. Top 75 shared hotspots in each genotype in nucleus Table S1. Library Information

Top 75 shared hotspots in (A) nucleus of WT BY4742 (libraries: FS106, FS142, FS213, FS214, FS215, FS53), (B) nucleus of rnh203-K46W BY4742 (libraries: FS325, FS334, FS335, FS336, FS337), (C) nucleus of rnh201-G42S (libraries: FS208, FS210, FS328) and (D) nucleus of rnh201△ BY4742 (libraries: FS109, FS138, FS322). In addition to the chromosomal coordinates, rNMP sequence context of the coordinate (3 nucleotides up/downstream from rNMPs), the mean Enrichment Factor and number of libraries containing rNMP and genome annotation are provided for each chromosomal coordinate of rNMP incorporation. (E) Composition of shared common hotspots in each genotype.

Table S5. Top 25 shared hotspots in each genotype in mitochondria

Top 25 shared hotspots in (A) mitochondria of WT BY4742 (libraries: FS106, FS142, FS213, FS214, FS215, FS53), (B) mitochondria of rnh203-K46W BY4742 (libraries: FS325, FS334, FS335, FS336, FS337), (C) mitochondria of rnh201-G42S (libraries: FS208, FS210, FS328) and (D) mitochondria of rnh201△ BY4742 (libraries: FS109, FS138, FS322). In addition to the chromosomal coordinates, rNMP sequence context of the coordinate (3 nucleotides up/downstream from rNMPs), the mean Enrichment Factor and number of libraries containing rNMP and genome annotation are provided for each chromosomal coordinate of rNMP incorporation. (E) Composition of shared common hotspots in each genotype

Table S6. P-values of preference using one sided Mann Whitney U test for normalized frequency values in nuclear genome heatmaps.

P-values of one-sided Mann-Whitney *U* test for nucleus normalized frequency values of (A) rNMP base (rA, rC, rG and rU) and dinucleotide combinations upstream NR (B) and downstream RN (C) of rNMP base (A, C, G and T for rA, rC, rG and rU). The test was conducted for all libraries and libraries present in each genotype against the expected frequency value of 0.25. P-value less than 0.05 (significant) are highlighted in red text.

Table S7. P-values of preference using one sided Mann Whitney U test for normalized frequency values in mitochondrial genome heatmaps.

P-values of one-sided Mann-Whitney *U* test for mitochondrial normalized frequency values of (A) rNMP base (rA, rC, rG and rU) and dinucleotide combinations upstream NR (B) and downstream RN (C) of rNMP base (A, C, G and T for rA, rC, rG and rU). The test was conducted for all libraries and libraries present in each genotype against the expected frequency value of 0.25. P-value less than 0.05 (significant) are highlighted in red text.

Table S8. P-values of genotype differences using two-sided Mann-Whitney *U* test for normalized frequency values in nuclear genome heatmaps.

P-values of two-sided Mann-Whitney *U* test for nuclear normalized frequency values of (A) rNMP base (rA, rC, rG and rU) and dinucleotide combinations upstream NR (B) and downstream RN (C) of rNMP base (A, C, G and T for rA, rC, rG and rU) for each genotype combinations. P-value less than 0.05 (significant) are highlighted in red text.

Table S9. P-values of genotype differences using two sided Mann-Whitney *U* test for normalized frequency values in mitochondrial genome heatmaps.

P-values of two-sided Mann-Whitney *U* test for mitochondrial normalized frequency values of (A) rNMP base (rA, rC, rG and rU) and dinucleotide combinations upstream NR (B) and downstream RN (C) of rNMP base (A, C, G and T for rA, rC, rG and rU) for each genotype combinations. P-value less than 0.05 (significant) are highlighted in red text.

## RESOURCE AVAILABILITY

### Lead contact

Further information and requests for resources and reagents should be directed to and will be fulfilled by the Lead Contact, Francesca Storici.

### Materials Availability

All unique/stable reagents generated in this study are available from the Lead contact.

### Data and code availability

The authors declare that the data supporting the findings of this study are available within the paper and its supplementary information files. The datasets generated during this study are available at NCBI’s Gene Expression Omnibus (GEO) repository under <u>GSE240399</u>. All data generated in this study are available from the Lead contact.

The following tools for analysis of data are available for download at GitHub under GNU GPL v3.0 license.

### Experimental Model and Study Participant Details

*Delitto Perfetto* System ^48^ was used for creating mutant strain using the background BY4742 (strain). The experimental method for analysis of single base resolution of ribonucleotide embedment used is ribose-seq^22,23^. Analysis used for sequencing data analysis of ribose-seq is using Ribose-map Toolkit^24,25^paired with modified shared or common hotspot analysis previously reported^49^ and nucleotide heatmaps using RibosePreferenceAnalysis software^50^ .

## METHOD DETAILS

### Multiple sequencing alignment of RNase H2 protein sequences

To observe the conserved domains between RNase H2 subunits of human and yeast, we used the amino acid sequences of RNase H2 catalytic subunit H2A, and accessory subunits H2B and H2C from *H. sapiens* and *S. cerevisiae*^9,32,51^. We performed multiple sequence alignment using Clustal Omega^52–54^ to predict conserved domains between all three subunits of RNase H2 enzyme complex in human and yeast. We mapped the amino acids involved in the catalytic activity^55^, conserved domains^32^ and yeast ortholog mutations previously reported^47^.

### Yeast strains and manipulations

We used haploid *Saccharomyces cerevisiae* strains from the BY4742 background. Strain BY4742 is a derivative of S288C, was used in the *S. cerevisiae* gene disruption project, and have *his3*Δ1 *leu2*Δ0 *lys2*Δ0 *ura3*Δ0 genotype with MATα mating type^56^. Standard genetic and molecular biology methods were used for growth, gene disruption, isolation of mutants, selection, genome engineering, colony PCR, and sequence analysis of genomic DNA^48,57,58^. Other genotypes were derived from KK-1^16^ by using the *delitto perfetto* method^48^ to generate *rnh201*-G42S (GGC->AGC codon change) and *rnh203*-K46W (AAG -> TGG codon change) to result in yeast orthologs of AGS-ortholog mutants. The *rnh201*-null genotype was derived using a previously reported method^16^. All mutations were confirmed by sequence analysis of PCR products obtained from amplification of a DNA region surrounding the specific mutation. The primers used for strain construction are listed in Table S2.

### Ribose-seq and library preparation

The analyzed ribose-seq libraries were constructed, following the previous protocols with a few modifications^18, 22, 23^. Three or more ribose-seq libraries were prepared for each genotype of the yeast strains using up to three different sets of restriction enzymes (RE1: DraI, EcoRV, and SspI; RE2: AluI, DraI, EcoRV, and SspI; and RE3: HaeIII and RsaI). This strategy allowed us (i) to verify that the conclusions taken from our analyses of ribose-seq data are not influenced by a particular set of restriction enzymes used to fragment the DNA extracted from the different yeast genotypes, and (ii) to further confirm reproducibility of the results. All ribose-seq libraries have a specific barcode within the sequence of the Unique Molecular Identifier (UMI) to distinguish the libraries from each other in the sequencing run and eliminate PCR duplicates^26^. All commercially available enzymes in the ribose-seq protocol were used according to the manufacturer’s instructions. In all DNA purification steps, nuclease free water was used to elute DNA. *S. cerevisiae* cells were inoculated in 150 mL of a YPD liquid medium containing yeast extract, peptone, and 2% (wt/vol) dextrose and incubated in a shaking incubator for 2 days to reach stationary phase. Genomic DNA was extracted using Qiagen Genomic DNA protocol “Preparation of Yeast Samples”. Successively, 40 μg of yeast genomic DNA was fragmented using restriction enzymes to produce blunt-ended fragments with an average size of 450 base pairs (bp) in length. Multiple sets of restriction enzymes were used for different library preparations, as shown in Table S1. The fragmented DNA was then purified by QIAquick PCR Purification Kit (QIAGEN). The purified DNA fragments were tailed with dATP (New England Biolabs, NEB) by using Klenow Fragment (3′→5′ exo-) (NEB) for 30 min at 37 °C and purified by QIAquick PCR Purification Kit. Following dA-tailing and purification, the DNA fragments were annealed with a partially double-stranded adapter (Adapter.L1∼L8 with Adapter.S, Table S<u>2</u>) by T4 DNA ligase (NEB) for overnight incubation at 16 °C. After that, the adapter-ligated products were purified using HighPrep PCR (MAGBIO). The annealed fragments were treated with a final concentration of 0.3 M NaOH for 2 h at 55 °C to denature the DNA strands, and to cleave at the rNMP sites resulting in 2′,3′-cyclic phosphate, 3’-phosphate, or 2′-phosphate termini. This was followed with neutralization using 2 M HCl and purification using HighPrep RNA Elite (MAGBIO). All the successive purification steps were performed using HighPrep RNA Elite (MAGBIO). The single-stranded DNA fragments were initially incubated at 95 °C for 3 min and then on ice for 2 min to fully denature any secondary structures of DNA. incubated with a final concentration of 1 μM *Arabidopsis thaliana* tRNA ligase (AtRNL), 50 mM Tris-HCl pH 7.5, 40 mM NaCl, 5 mM MgCl2, 1 mM DTT, and 300 μM ATP in a volume of 20 μL for 1 h at 30 °C and followed by purification. AtRNL ligates the 2′-phosphate ends of rNMP-terminated ssDNA fragment to its opposite 5′-phosphate end, which results in a circular ssDNA.

The fragments were then treated with T5 Exonuclease (NEB) in a volume of 50 μL for 30 min at 37 °C to degrade all remaining linear ssDNA fragments. After purification, the circular fragments were incubated with a final concentration of 1 μM 2′-phosphotransferase (Tpt1), 20 mM Tris-HCl pH 7.5, 5 mM MgCl2, 0.1 mM DTT, 0.4% Triton X-100, and 10 mM NAD^+^ in a volume of 40 μL for 1 h at 30 °C to remove the 2′- phosphate presenting at the ligation junction. After Tpt1 treatment and purification, the circular fragments were amplified by two rounds of polymerase chain reaction (PCR) with Q5-High Fidelity polymerase (NEB). Both PCR rounds begin with an initial denaturation at 98 °C for 30 s. Denaturation at 98 °C for 10 s, primer annealing at 65 °C for 30 s, and DNA extension at 72 °C for 30 s are then performed. These three steps are repeated for 6 cycles in the first PCR round, and for 11 cycles in the second PCR round. Successively, there is a final extension reaction at 72 °C for 2 min for both PCRs.

The first round of PCR was performed to amplify and introduce the adapter sequences of Illumina TruSeq CD index primers. The primers PCR.1 and PCR.2 used for the first round were the same for all libraries. The second round of PCR was performed to attach specific indexes i7 and i5 for each library. The sequences of PCR primers and indexes can be found in Table S2. The amplified DNA fragments were then performed with a double-sided size selection to select specific sizes of DNA between 250 bp and 650 bp using HighPrep PCR (MAGBIO). The resulting ribose-seq libraries were quantified with Qubit and Bioanalyzer. The prepared libraries were sequenced on the Illumina HiSeq X ten and NextSeq 500 platforms at Admera Health. For this study, we built 4 libraries for analysis of wild-type RNase H2, five libraries for *rnh203*-K46W, three libraries for *rnh201*-G42S, and two libraries for *rnh201*△. For controls, we also included two wild-type RNase H2 (FS106 and FS142)^18^ and one *rnh201*△ libraries (FS138)^18^ that were previously published.

### Preliminary data processing and alignment

For the ribose-seq libraries, the sequencing reads consist of an eight-nucleotide UMI, a three-nucleotide molecular barcode, the tagged nucleotide (the nucleotide tagged during ribose-seq from which the position of the rNMP is determined), and the sequence directly downstream from the tagged nucleotide. The UMI corresponds to sequence position 1–6 and 10–11, the molecular barcode corresponds to position 7–9, the tagged nucleotide corresponds to position 12, and the tagged nucleotides downstream of the sequence corresponds to positions 13+ of the raw FASTQ sequences. The rNMP is the reverse complement of the tagged nucleotide. Before aligning the sequencing reads to the reference genome, the reads were trimmed based on sequencing quality and the custom ribose-seq adaptor sequence using cutadapt 1.16 (-q 15 -m 62 -a “AGTTGCGACACGGATCTATCA”). In addition, to ensure accurate alignment to the reference genome, reads containing fewer than 50 nucleotides of genomic DNA (those nucleotides located downstream from the tagged nucleotide) after trimming were discarded. Following quality control, the Alignment and Coordinate Modules of the Ribose-Map toolkit were used to process and analyze the reads^26^. The Alignment Module de-multiplexed the trimmed reads by the appropriate molecular barcode, aligned the reads to the reference genome (sacCer2) using Bowtie 2, and de- duplicated the aligned reads using UMI-tools, as also mentioned in previously published study about ribose-seq analysis on yeast strains^16^.. Based on the alignment results, the Coordinate Module filtered the reads to retain only those with a mapping quality score of at least 30 (probability of misalignment <0.001) and calculated the chromosomal coordinates and per-nucleotide counts of rNMPs.

Due to the efficient removal of rNMPs by RNase H2, nuclear libraries of wild-type cells generally had a much lower number of reads compared to the mitochondrial libraries of the same cells, and thus had a higher number of background reads that originated from the capture of restriction enzyme ends likely by residual activity of T4 DNA ligase. To improve the quality of analysis and filter out any background reads or artifacts, we filtered out those reads that showed restriction enzyme cleavage sites at rNMP incorporation sites, which removes any linear single strand DNA-only fragments being captured. We then also filter out rNMP mismatches between the nucleotide in reference genome versus sequencing reads. The final file generated is of BED format and all further analysis is performed this bed file after filtration of restriction enzyme site and mismatched rNMPs.

### Ribonucleotide percentages, composition, and sequence context analysis

Since each library can have different coverage, we used percentage of rNMPs in nuclear and mitochondrial DNA to determine the rate of incorporation in difference genotypes. We then visualized these percentages using horizontal bar graphs generated using customized R scripts.

The Composition Module of Ribose-Map was used to obtain raw and normalized percentages of each rNMP base (A, C, G, U/T) in nuclear and mitochondrial DNA of all genotypes. The normalized percentages are the raw rNMP counts normalized on the nucleotide counts in the corresponding (nuclear or mitochondrial) reference genome. The Sequence Module of Ribose-Map was used to generate nucleotide sequence context plots encoded using the R-ggplot package. The 1 % of the most abundant rNMP sites in the genome of each library were selected as hotspots, and the consensus sequence around these hotspots was visualized using customized R scripts and plotted using ggseqlogo^59^.

### Ribonucleotide count normalization and mapping shared hotspots

DNA extracted from the yeast cells for ribose-seq was also sequenced using Illumina HiSeq X Ten Platform. The fastq reads were trimmed for any residual Illumina adapters and filtered to keep reads above the quality threshold of 15 and length 50 bp. These trimmed reads were then aligned using bowtie2.0 with sacCer2 genome to get an alignment file in SAM (Sequence Alignment Map) format. The SAM file was then sorted and indexed to a BAM (Binary Alignment Map) file.

Coverage per base pair (*C*_*b*_) and mean genomic coverage per base pair (*C*_*gb*_) was calculated using SAMtools. Relative base coverage (*C*_*Rl*_) of each base (*b*) in the genome is calculated as follows:

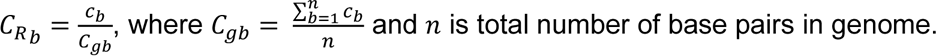

This relative coverage per base (*C*_*Rb*_) is further used to calculate rNMPs per base coverage (*R*_*Cb*_) by normalizing the rNMP counts per base pair (*R*_*b*_) using the following formula:

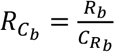

We calculate *R*_*Cb*_ for each base pair in each Library using respective genotype coverage information. We also calculate Enrichment Factor *EF*_*Cb*_ using rNMPs per base coverage (*R*_*Cb*_) for each base in each library.

The enrichment factor is calculated relative to the length of genome and region of interest, region here being each base pair.

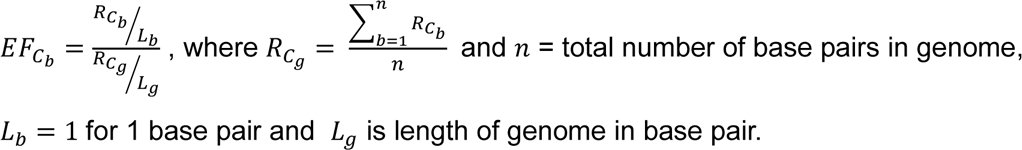

*R*_*Cb*_ rounded to the closest integer ⌊*R*_*Cb*_⌉ in each Library is used to calculate the presence of rNMPs in each library of every genotype. Top shared hotspot locations in each genotype are determined based on highest mean Enrichment Factor and recurrence of rNMPs in a genomic location in at least 2 libraries of each genotype.

Nucleotide-frequency analysis and visualization using heatmaps.

To generate the mononucleotide heatmaps, in every nuclear and mitochondrial ribose-seq library, the frequency of each type of rNMP (FN: FA, FC, FG, or FU) was calculated as a ratio of percentage of each rNMP base divided by the percentage of each dNTP base in the reference genome. Normalized frequency for each rNMP base was calculated using the frequency of the respective rNMP base divided by the sum of frequencies of each rNMP base^16,60^. The sum of all normalized frequencies in the mononucleotide heatmaps sums up to 1.

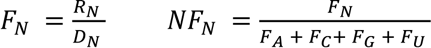

RN -> percentage of ribo-nucleotides where N ∈ {A, C, G, U}

DN -> percentage of di-nucleotides in reference genome where N ∈ {A, C, G, T}

For dinucleotide heatmaps, normalized frequency was calculated for each deoxyribonucleotide immediately upstream or downstream of rNMP incorporation site for each base such that the sum of NR (upstream) or RN (downstream) frequencies for each R would sum upto 1^16,60^.

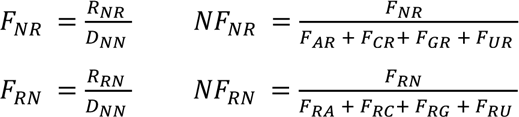

RNR -> percentage of dinucleotide upstream of ribo-nucleotides where N ∈ {A, C, G, T} and R ∈ {A, C, G, U}

RRN -> percentage of dinucleotide downstream of ribo-nucleotides where R ∈ {A, C, G, U} and N ∈ {A, C, G, T}

DNN -> percentage of di-nucleotides in reference genome where N ∈ {A, C, G, T}

### ARS region leading and lagging strand heatmap analysis

ARS annotations of OriDB were downloaded and transferred to the sacCer2 reference genome with Liftover software. Only confirmed ARSs were included (n = 410^61^. Among them, 276 ARSs have a known firing time^62^ . They were divided into two halves, early-firing ARSs (T ≤ 24.7 min, n = 139) and late-firing ARSs (T > 24.7 min, n = 137). The upstream and downstream 15-kb regions are considered as the flanks of an ARS. The 5′-upstream flanks of ARSs for both the Watson and Crick strands correspond to the lagging strand, and the 3′-downstream flanks of ARSs correspond to the leading strand. If two ARSs are close to each other and the distance between them is smaller than 30 kb, the position of the collision point of the corresponding converging replication forks is calculated with their firing times and the average fork moving speed of 1.6 kb/min^63^ . Bedtools^64,65^ was used to get rNMPs overlapping in 4 -10 kb of leading and lagging strand from early firing ARS to generate dinucleotide frequency heatmaps for each rNMP and normalized on the dinucleotide frequencies found in respective strand.

## QUANTIFICATION AND STATISTICAL ANALYSIS

We used a one-sided Mann-Whitney *U* test to compare frequency results of heatmap data with expected frequency of 0.25 for each rNMP base and dinucleotide combination, in nuclear and mitochondrial DNA of each genotype library. We also did two-sided Mann-Whitney U test to compare frequencies of AGS mutants or *rnh201*△ mutant with wild-type frequencies. *P*-values of these tests are indicated in Table S6- 9.

