## Supplementary Fig Legends for "Distinct features of ribonucleotides within genomic DNA in Aicardi-Goutières syndrome (AGS)-ortholog mutants of *Saccharomyces cerevisiae*"

**SUPPLEMENTARY FIGURE LEGENDS**

**Figure S1. Raw ribonucleotide base percentages in BY4742 AGS mutant genotype libraries, related to Figure 2B-C**

Bar graphs showing the mean value and standard error of each ribonucleotide base percentages of rNMPs found in (A) nuclear and (B) mitochondrial genome of rNMP libraires for the indicated genotypes. The percentages are directly calculated from counts of each Ribonucleotide base (R) by the total Ribonucleotides found in nuclear and mitochondrial genome respectively in each library. The percentages are mentioned in the Supplementary Table. On the right is the legend showing colors used for each ribonucleotide base. Red, A; blue, C; yellow, G; and green, U.**­**

**Figure S2. Sequence frequency plots flanking ribonucleotide incorporation site for all and each ribonucleotide base for all ribose-seq libraries, related to Figure 2D-E**

Sequence context +/- 100 nucleotides (zoomed out) and +/- 5 nucleotides (zoomed in) plots of normalized nucleotide frequencies from the site of rNMP presence, which is indicated by the 0 position, for all rNMPs found in nuclear and (E) mitochondrial DNA, respectively. relative to mapped positions of sequences from nuclear (A-B) and mitochondrial (C-D) ribose-seq libraries. The frequency of each nucleotide is normalized to the frequency of the corresponding nucleotide present in the nuclear or mitochondrial reference genome. The plots shown are for one sample library of each genotype with the library and genotype indicated on left of the panels. Negative x-axis labels denote upstream nucleotide position whereas positive x-axis labels denote downstream nucleotide position from the site of incorporation. The Y-axis denotes Log 2 scale of nucleotide frequencies normalized to the corresponding frequencies of sacCer2 nuclear or mitochondrial genome. The colors and shapes of each nucleotide base is shown as; Red square, A; blue circle, C; orange triangle, G; and green rhombus, U. The ribonucleotide base for the displayed data is indicated on top of each plot. Combined plots show all rNMPs and sequence context around normalized based of all dNMPs in corresponding genome, whereas A, C, G, U plots show only the corresponding rNMP in the 0^th^ positions and sequence context frequencies normalized to dA, dC, dG and dT respectively in the corresponding genome.

**Figure S3. Sequence logo plots 3 bases around for top 1 percentile most abundant ribonucleotide incorporation sites (hotspots) in each library, related to Figure 2F-G**

Sequence logo plots flanking +/- 3 nt from the rNMP position (0) of top 1 percentile locations with the highest rNMP counts observed in (A) nuclear and (B) mitochondrial DNA. Negative values on x-axis represent nucleotide conservation upstream and positive denote conservation downstream for the site of incorporation. The *y*-axis shows the level of sequence conservation, represented in bits. The library name and genotype for the displayed data are indicated on top of each plot. The counts of rNMPs in the top 1 percentile position and total rNMP counts in each library and genome are represented at the bottom of each plot.

**Figure S4. Sequence logo plots 3 bases around and normalized percentages of most abundant ribonucleotide incorporation sites (hotspots) in each genotype, related to figure 3.**

Sequence logo plots flanking +/- 3 nt of top 75 count, 2 and 5 percentile of most abundant shared hotspot locations in (A) nucleus and top 25 count, 2 and 5 percentile of most abundant shared hotspot locations in (B) mitochondria. 0^th^ position denotes rNMP incorporation site, negative and positive values on x-axis represent nucleotide conservation upstream and downstream for the site of incorporation respectively. . The *y*-axis shows the level of sequence conservation, represented in bits. The counts of 1 and 2 percentile of shared hotspots in each genotype and genome are represented on the bottom of each plot. The represented annotations for rA, rC, rG and rU are shown in red, blue, yellow, and green color letters respectively. The shared hotspots are selected based on recurrence in at least 2 libraries in each genotype and highest rNMP Enrichment Factor(see Methods) normalized on relative genomic coverage of each genotype. The figure also shows normalized percentages of top 75 shared hotspots in the nucleus (C) and top 25 shared hotspots in the mitochondria (D). These percentages are normalized on dNMP percentages in corresponding genomes.

**Figure S6. Downstream dinucleotides(NR) preference in Leading and Lagging strands of early firing Autonomous Replicating Sequences (ARS) of AGS mutants, related to Figure 5.**

Dinucleotide RN (rA, rC, rG, and rU with the downstream deoxyribonucleotide with base A, C, G, or T) Heatmap analysis to reveal preferences on the (A) leading and (B) lagging strands for wild-type, *rnh201*△ , and AGS mutants 4,000 -10,000 nt from the early-firing ARSs. The formulas used to calculate these normalized frequencies are shown and explained in Methods. Each column of the heatmap shows results of a specific ribose-seq library. Libraries with more than 400 rNMPs in leading and lagging strand are displayed in the Heatmaps. Each library name is indicated underneath each column of the heatmap with its corresponding restriction enzyme (RE) set used. The ribose-seq libraries of the same genotype are also grouped together by curly brackets and separated by thick vertical blue lines. Each row shows results obtained for ribonucleotide or dinucleotide combination.  The actual percentages of rNMP (rA, rC, rG, and rU) dinucleotides of fixed base A, C, G, or T for the indicated base combinations (AA, AC, AG, and AT; CA, CC, CG, and CT; GA, GC, GG, and GT; and UA, UC, UG, and UT) present in 4,000 – 10,000 nts of leading and lagging strands are shown to the left of the heatmaps. The observed % of dinucleotides with rNMPs with base A, C, G, or U were divided by the actual % of each rNMP and dinucleotide with fixed base A, C, G, or T in 4,000 – 10,000 nts of leading and lagging strands. The bar to the right shows how different frequency values are represented as different colors: white for 0.25; light red to red for 0.25 to 0.5–1, and dark blue to light blue for 0.25 to 0.
