## Supplementary material for "Distinct features of ribonucleotides within genomic DNA in Aicardi-Goutières syndrome (AGS)-ortholog mutants of *Saccharomyces cerevisiae*": Figure S

### Figure S1. Ribonucleotide base percentages without normalization

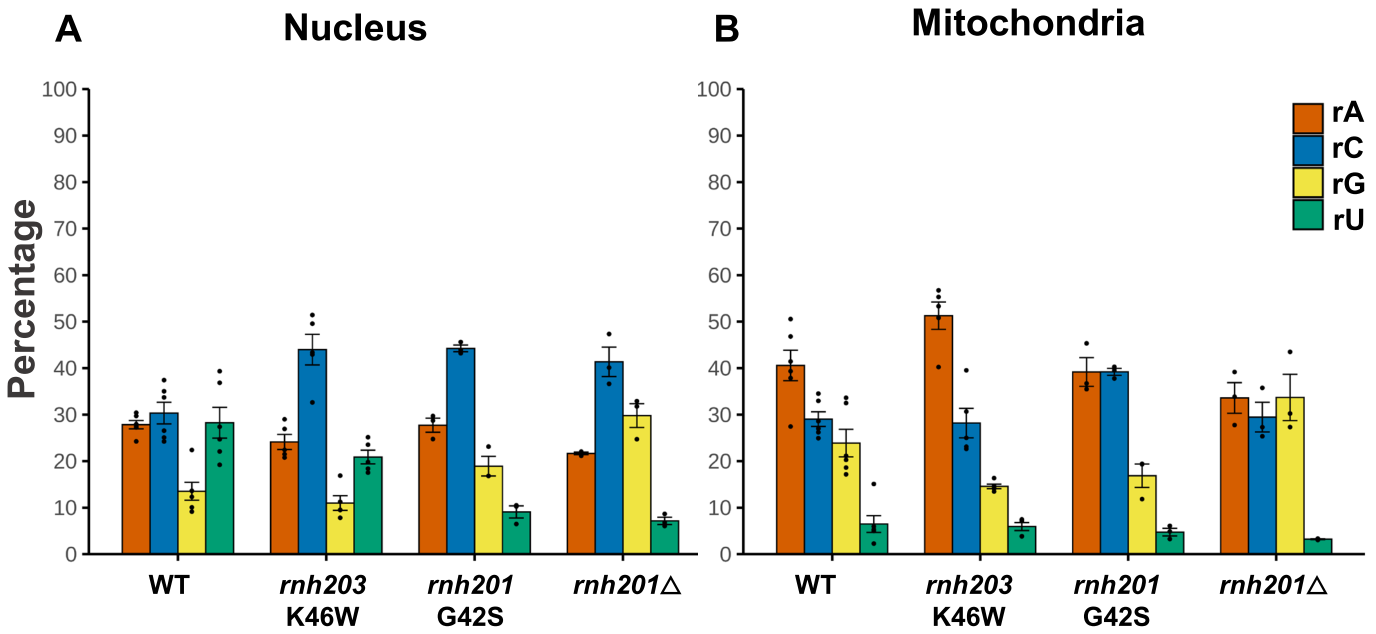

### Figure S2A. Sequence frequency plots around rNMPs in nucleus

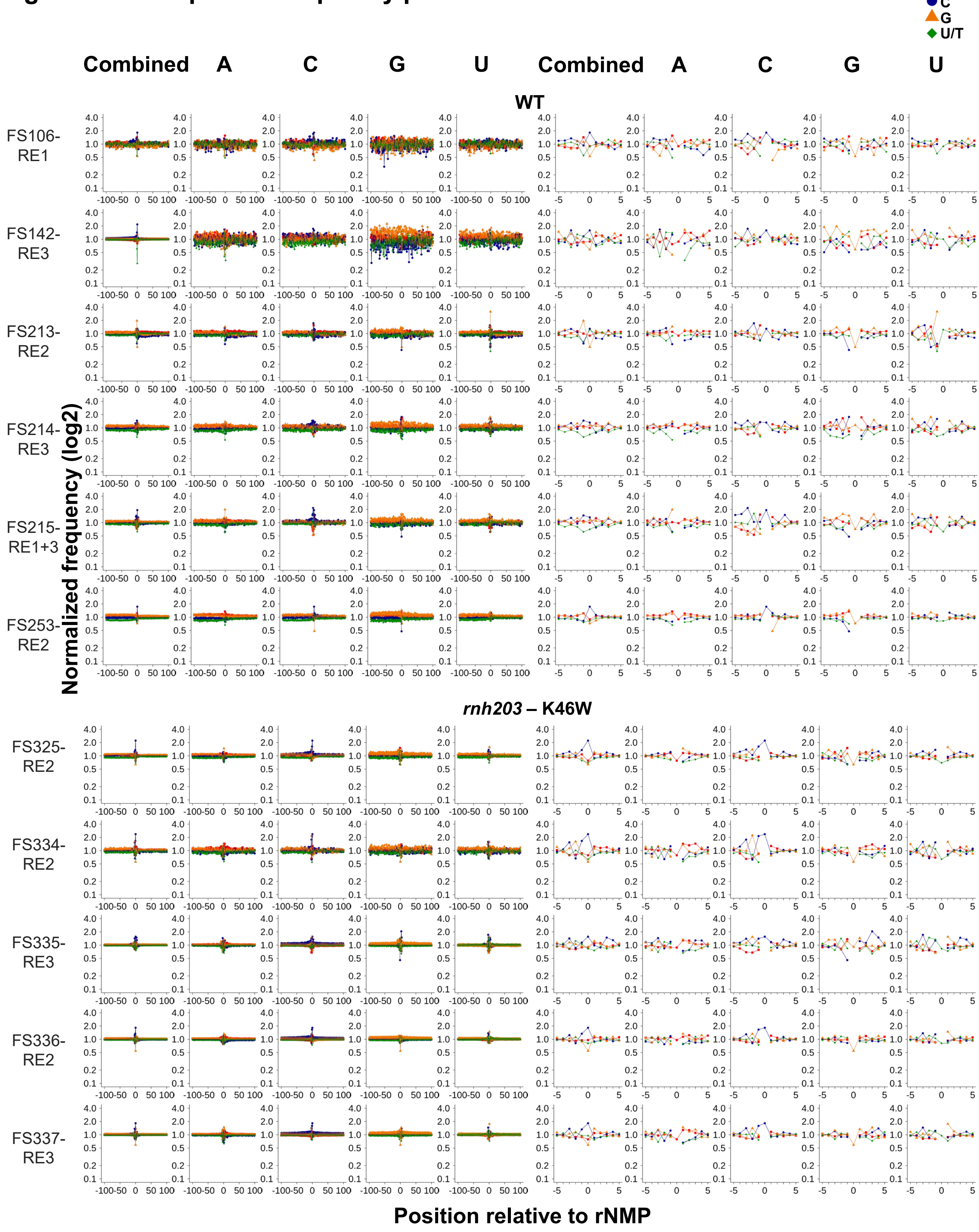

### Figure S2B. Sequence frequency plots around rNMPs in nucleus

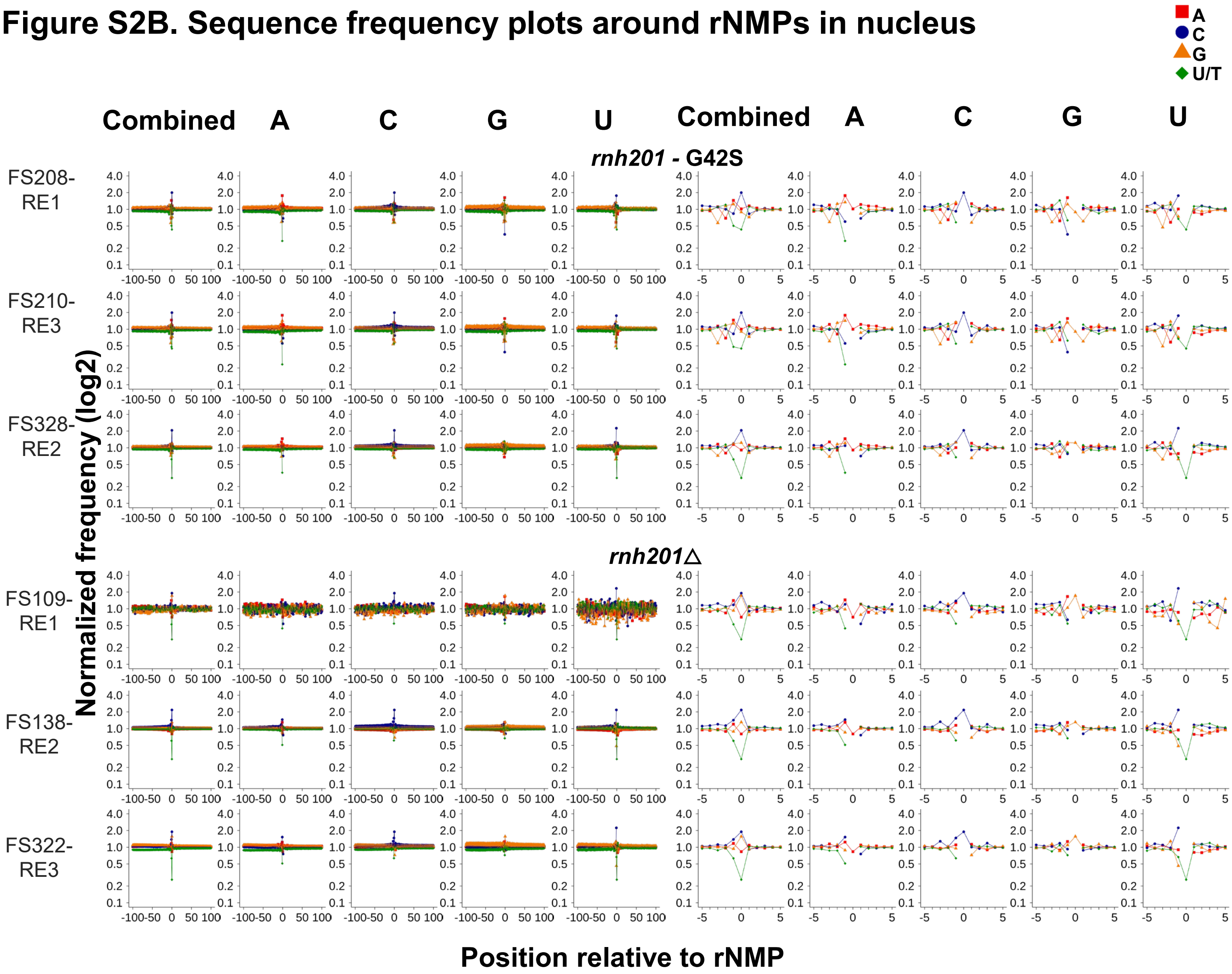

### Figure S2C. Sequence frequency plots around rNMPs in mitochondria

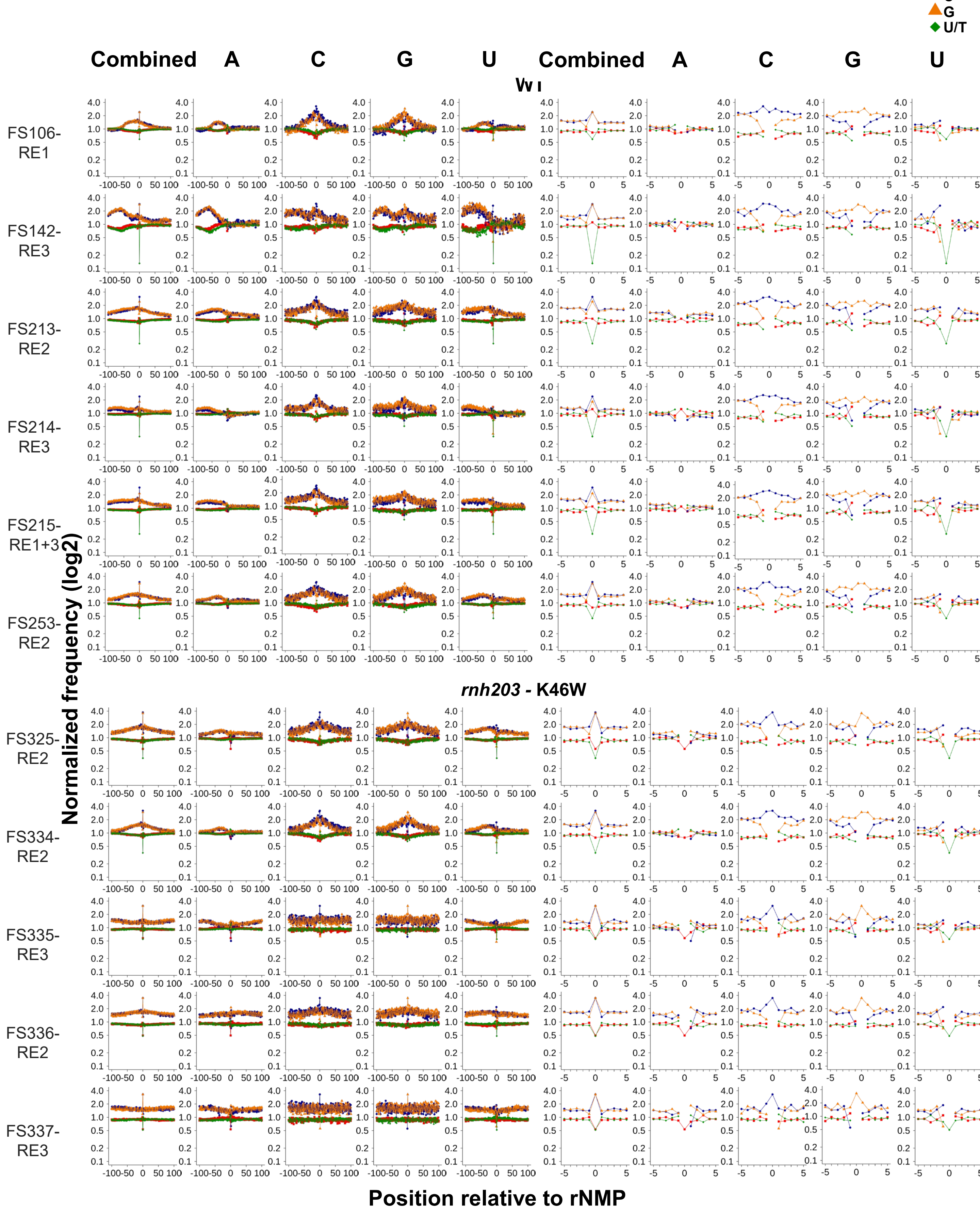

### Figure S2D. Sequence frequency plots around rNMPs in mitochondria

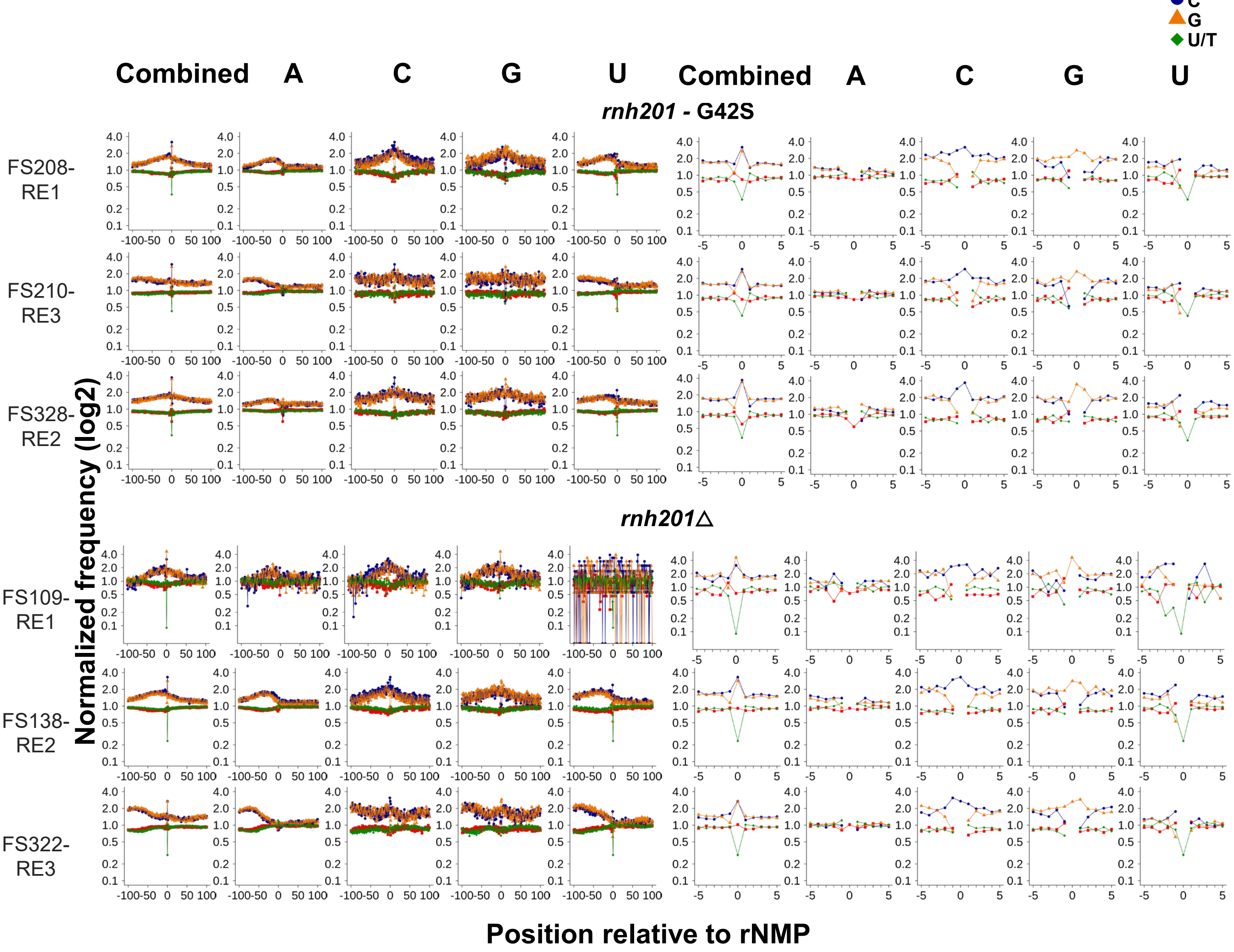

**Figure S3. Sequence logo plots for 1 percentile highly abundant rNMP sites**

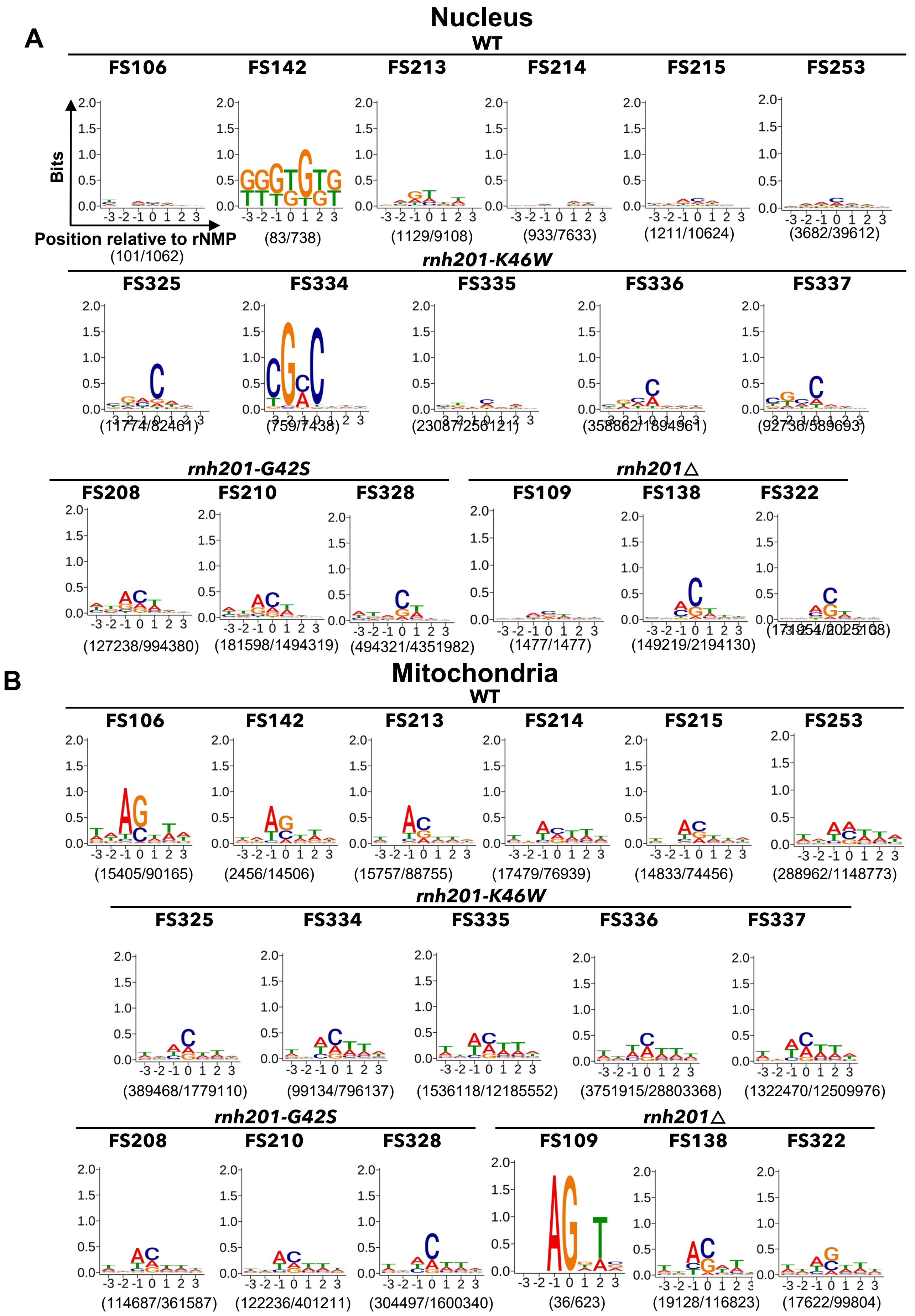

**Figure S4. Sequence logo plots and compositions for shared hotspots**

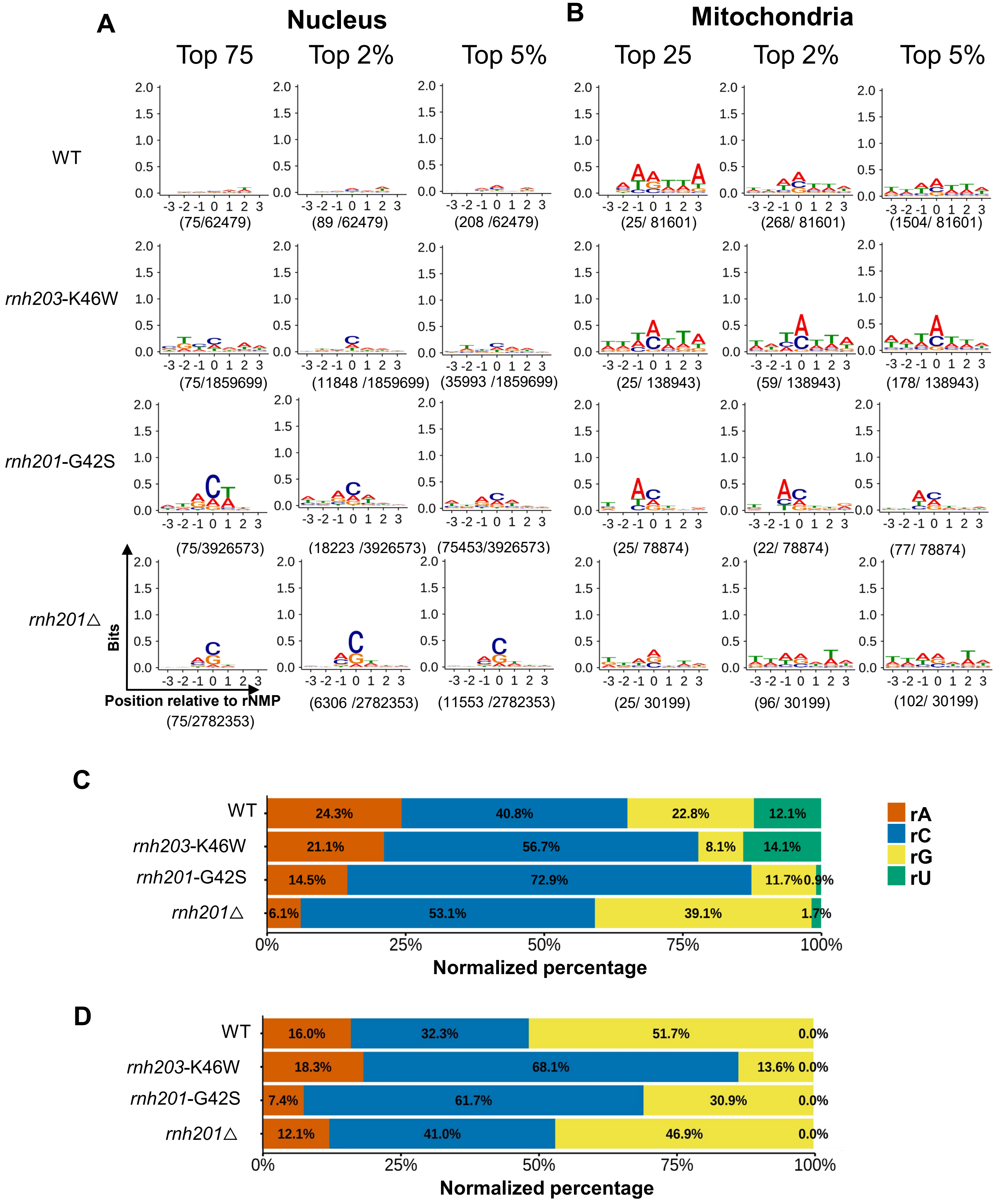

### Figure S5. Dinucleotide preferences in nucleus and mitochondria

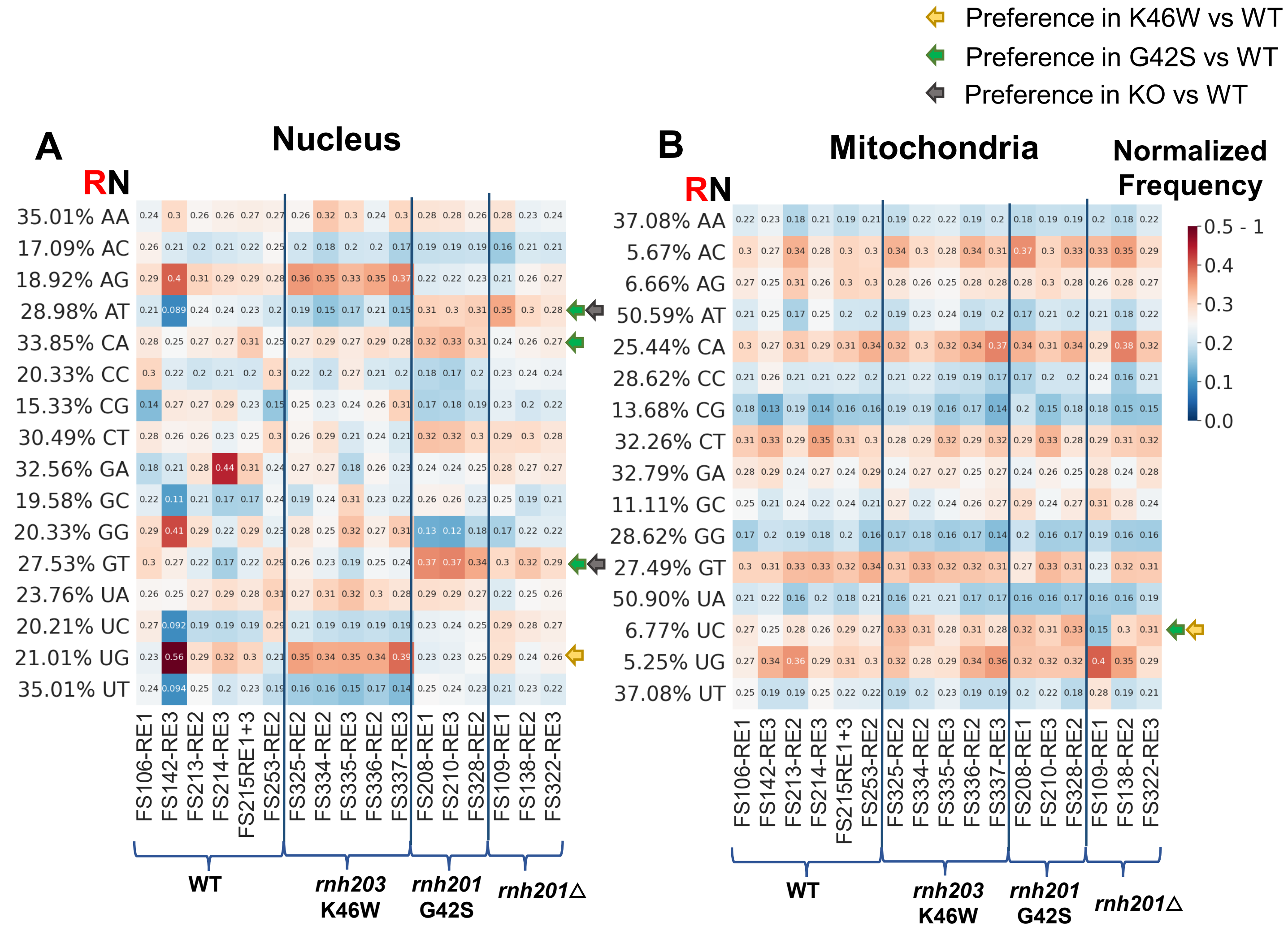

### Figure S6. Di-nucleotide preferences in 4-10 kb of DNA replication strands

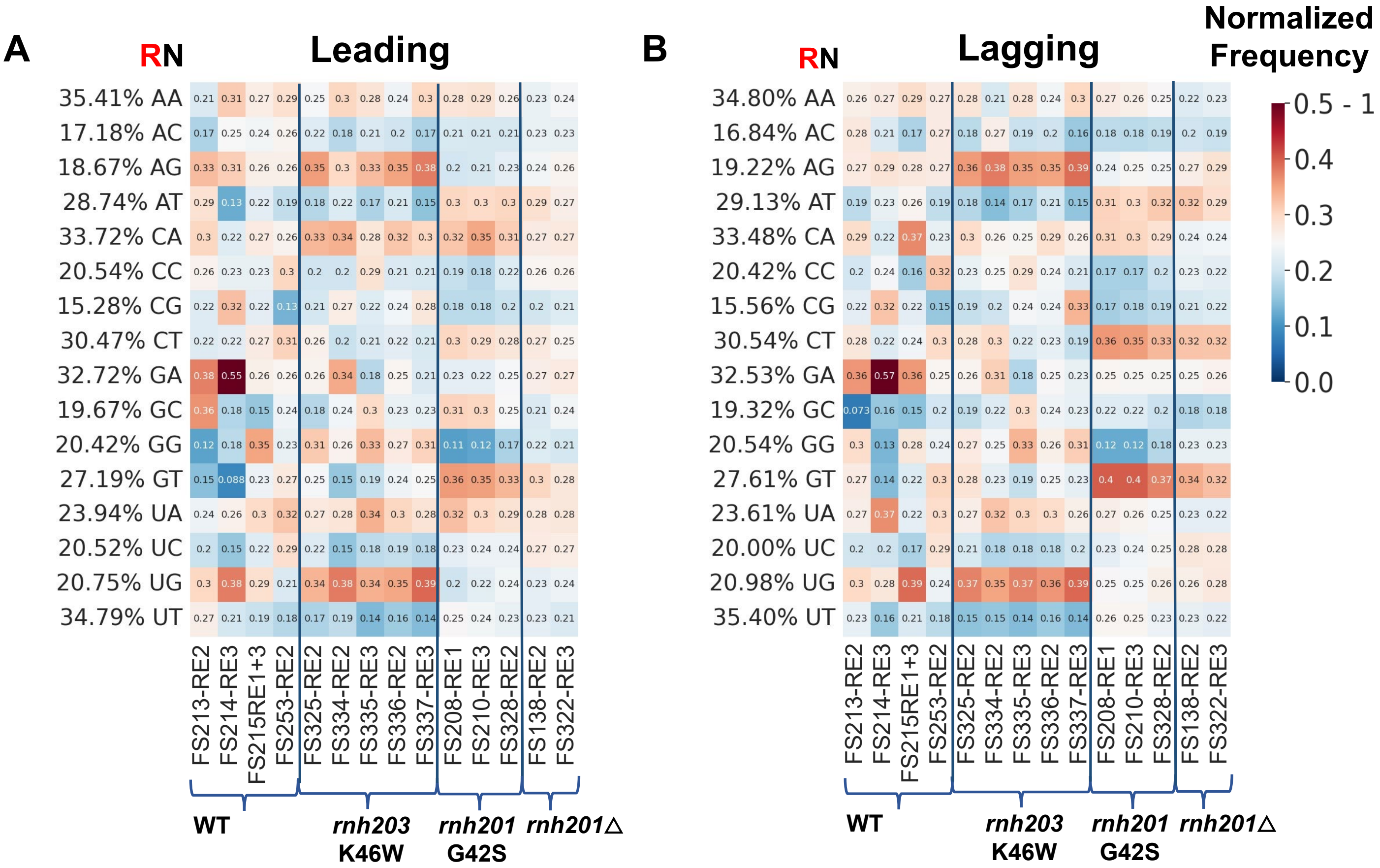
