## Supplementary material for "Distinct features of ribonucleotides within genomic DNA in Aicardi-Goutières syndrome (AGS)-ortholog mutants of *Saccharomyces cerevisiae*": Table S2

**Table S2. Oligonucleotides used in this study.**

| Name | Size | Sequence |
| --- | --- | --- |
| Adaptor.L1 * | 65 | 5’ P-NNC CGN NNN NNA GAT CGG AAG AGC GTC GTG TAG GGA AAG AGT GTT GAT AGA TCC GTG TCG CAA C*T |
| Adaptor.L2 * | 65 | 5’ P-NNT GAN NNN NNA GAT CGG AAG AGC GTC GTG TAG GGA AAG AGT GTT GAT AGA TCC GTG TCG CAA C*T |
| Adaptor.L3 * | 65 | 5’ P-NNG ACN NNN NNA GAT CGG AAG AGC GTC GTG TAG GGA AAG AGT GTT GAT AGA TCC GTG TCG CAA C*T |
| Adaptor.L5 * | 65 | 5’ P-NNG CTN NNN NNA GAT CGG AAG AGC GTC GTG TAG GGA AAG AGT GTT GAT AGA TCC GTG TCG CAA C*T |
| Adaptor.L6 * | 65 | 5’ P-NNA GCN NNN NNA GAT CGG AAG AGC GTC GTG TAG GGA AAG AGT GTT GAT AGA TCC GTG TCG CAA C*T |
| Adaptor.L8 * | 65 | 5’ P-NNT GTN NNN NNA GAT CGG AAG AGC GTC GTG TAG GGA AAG AGT GTT GAT AGA TCC GTG TCG CAA C*T |
| Adaptor.S * | 25 | 5’ P-GTT GCG ACA CGG ATC TAT CAA CAC T -Am 3’ |
| PCR.1 | 54 | 5’ GTG ACT GGA GTT CAG ACG TGT GCT CTT CCG ATC TTG ATA GAT CCG TGT CGC AAC |
| PCR.2 | 20 | 5’ ACA CTC TTT CCC TAC ACG AC |
| PCR.701 | 53 | 5’ CAA GCA GAA GAC GGC ATA CGA GAT **CGA GTA AT**G TGA CTG GAG TTC AGA CGT GT |
| PCR.702 | 53 | 5’ CAA GCA GAA GAC GGC ATA CGA GAT **TCT CCG GA**G TGA CTG GAG TTC AGA CGT GT |
| PCR.705 | 53 | 5' CAA GCA GAA GAC GGC ATA CGA GAT **TTC TGA AT**G TGA CTG GAG TTC AGA CGT GT |
| PCR.712 | 53 | 5' CAA GCA GAA GAC GGC ATA CGA GAT **CTA TCG CT**G TGA CTG GAG TTC AGA CGT GT |
| PCR.501 | 57 | 5’ AAT GAT ACG GCG ACC GAG ATC TAC AC**T ATA GCC T**AC ACT CTT TCC CTA CAC GAC |
| PCR.502 | 57 | 5’ AAT GAT ACG GCG ACC GAG ATC TAC AC**A TAG AGG C**AC ACT CTT TCC CTA CAC GAC |
| PCR.503 | 57 | 5’ AAT GAT ACG GCG ACC GAG ATC TAC AC**C CTA TCC T**AC ACT CTT TCC CTA CAC GAC |
| PCR.504 | 57 | 5’ AAT GAT ACG GCG ACC GAG ATC TAC AC**G GCT CTG A**AC ACT CTT TCC CTA CAC GAC |
| PCR.505 | 57 | 5’ AAT GAT ACG GCG ACC GAG ATC TAC AC**A GGC GAA G**AC ACT CTT TCC CTA CAC GAC |
| PCR.506 | 57 | 5’ AAT GAT ACG GCG ACC GAG ATC TAC AC**T AAT CTT A**AC ACT CTT TCC CTA CAC GAC |
| PCR.507 | 57 | 5’ AAT GAT ACG GCG ACC GAG ATC TAC AC**C AGG ACG T**AC ACT CTT TCC CTA CAC GAC |
| PCR.508 | 57 | 5’ AAT GAT ACG GCG ACC GAG ATC TAC AC**G TAC TGA C**AC ACT CTT TCC CTA CAC GAC |

Name, length, and sequence of oligonucleotides used in this study are presented. All bold letters in the PCR primers indicate the specific sequence of index used in sequencing. P and Am indicate end modifications of phosphate and amino modifier, respectively. All oligonucleotides were desalted, except those marked with an asterisk (*), which were HPLC purified. All oligonucleotides were synthesized by Integrated DNA Technologies.
